# Implication of the cellular factor CTCF in the regulation of Bovine Leukemia Virus latency and tridimensional chromatin organization

**DOI:** 10.1101/2021.08.28.457911

**Authors:** Anthony Rodari, Maxime Bellefroid, Mathilde Galais, Peter H.L. Krijger, Lorena Nestola, Estelle Plant, Erica S.M. Vos, Benoit Van Driessche, Caroline Vanhulle, Amina Ait Ammar, Angela Ciuffi, Wouter de Laat, Carine Van Lint

**Affiliations:** Service of Molecular Virology, Department of Molecular Biology (DBM), Université Libre de Bruxelles (ULB), Gosselies, 6041, Belgium; Oncode Institute, Hubrecht Institute-KNAW and University Medical Center Utrecht, Utrecht, 3584 CT, the Netherlands; Institute of Microbiology, Lausanne University Hospital, University of Lausanne, Lausanne, 1011, Switzerland

## Abstract

Bovine Leukemia Virus (BLV)-induced tumoral development is a multifactorial phenomenon which remains largely unelucidated. Here, we highlighted the critical role of the cellular CCCTC-binding factor (CTCF) both in the regulation of BLV transcriptional activities and in the deregulation of the tridimensional (3D) chromatin architecture surrounding the BLV integration site. We demonstrated the in vivo recruitment of CTCF to three conserved CTCF binding motifs along the BLV provirus. Next, we showed a critical role for CTCF in delimitating the epigenetic landscape along the BLV provirus as well as to repress the 5’Long Terminal Repeat (LTR) promoter activity, thereby contributing to viral latency, while favoring the 3’LTR promoter activity. Finally, we demonstrated that BLV integration deregulated host cellular 3D chromatin organization through the formation of abnormal viral/host chromatin loops. Altogether, our results highlight CTCF as a new critical effector of BLV transcriptional regulation and BLV-induced physiopathology.

## INTRODUCTION

Bovine leukemia virus (BLV) is a B-lymphotropic oncogenic deltaretrovirus infecting cattle. Infections are under control in Western Europe with the help of regulated sanitary rules whereas countries lacking these measures are still facing massive economical losses in the food and milk industries (1, 2). In addition to this economical issue, BLV shares common features with Human T-lymphotropic Virus 1 and 2 (HTLV-1 and -2), thereby constituting a convenient animal model to further study the HTLV-1-dependent tumorigenesis in humans (3, 4). Regarding BLV physiopathology, while the majority of BLV-infected animals remain asymptomatic lifelong, 30% of them will develop a persistent lymphocytosis and less than 5% will suffer from B-cell leukemia or lymphoma, termed enzootic bovine leukosis, leading to a rapid death of BLV-infected cattle (4). Remarkably, to further study BLV-induced oncogenic mechanisms, BLV can be inoculated experimentally to sheep, who have been demonstrated to be more sensitive to BLV-associated oncogenic properties with 95% of the infected animals developing B-cell leukemia or lymphoma after a shorter period of incubation compared to bovines (5, 6). Overall, a common feature of BLV infection is the viral latency, characterized by an absence of viremia (4, 7, 8), which probably enables the escape from the host immune system and ultimately tumor development (9). Mechanistically, we and others have demonstrated that this viral latency occurs through the repression of the RNA polymerase II-dependent (RNAPII) promoter located in the 5’long terminal repeat (5’LTR) by several mechanisms (10), including genetic mutations in important *cis*-regulatory regions (11, 12), epigenetic modifications (13–17), as well as the sequestration of host cellular transcription factors for which binding sites have been previously identified along the 5’LTR (18–25). However, despite the deep repression affecting the 5’LTR, we and others have discovered and characterized two additional promoter activities (26–29). Indeed, BLV encodes a highly expressed miRNA cluster, responsible for transcription of 10 viral miRNAs through a non-canonical process involving the RNA polymerase III (RNAPIII). In addition, the 3’LTR exhibits an important RNAPII-dependent promoter activity, responsible for the expression of 3 viral antisense transcripts (26, 29). Functionally, the BLV miRNAs and antisense transcripts could be responsible for tumor progression by deregulating the host transcriptome (30, 31) and producing viral/host chimeric transcripts (32), respectively, in infected animals. Altogether, these findings have provided new insights into alternative ways used by BLV to express parts of its genome, despite viral latency affecting the 5’LTR promoter activity, thereby bringing additional approaches to study BLV-induced leukemogenesis.

In the present report, we investigated the putative role of the cellular CCCTC-binding factor (CTCF) in the regulation of BLV RNAPII-dependent promoter activities through transcriptional and epigenetic mechanisms. We also investigated its implication in host gene expression deregulation through the formation of abnormal viral/host chromatin loops. Indeed, CTCF is a transcription factor ubiquitously expressed in all cell types and highly conserved across bilaterians. Functionally, CTCF has been demonstrated to play major roles in transcriptional and epigenetic regulation, mainly by organizing the cellular genome at the tridimensional (3D) level. CTCF-mediated regulations occurs through the formation of chromatin loops, in cooperation with the cohesin multiprotein complex. Besides “high-order” regulation, local and direct effects on gene expression have also been described, depicting CTCF as an important multifunctional protein (reviewed in (33–36)). Of note, CTCF has been demonstrated to be involved in the regulation of the infectious cycle of several DNA viruses such as Kaposi’s sarcoma-associated herpesvirus (KSHV), human papillomavirus (HPV) and Epstein-Barr virus (EBV) (37). Regarding retroviruses, CTCF has been shown to be recruited to a unique proviral binding site located in the regulatory region of the HTLV-1 provirus (38). However, the functional role of CTCF recruitment in HTLV-1 gene expression regulation as well as in HTLV-1-induced tumorigenesis remains incompletely understood (38–42). In the present manuscript, we demonstrate a critical role of CTCF as a regulator of BLV gene expression and possibly as a new determinant of BLV-induced leukemogenesis.

## MATERIAL AND METHODS

### Cell lines and primary cell samples

The ovine cell lines L267 (15) and YR2 (8) are clonal cell lines established from the T267 B-cell lymphoma and M395 B-cell leukemia, respectively, developed by a BLV-infected sheep injected with naked proviral DNA of an infectious BLV variant (6) displaying a wild-type sequence. These cell lines were maintained in Opti-MEM GlutaMAX medium (Life Technologies) supplemented with 10% FBS, 1 mM sodium pyruvate, 2 mM glutamine, non-essential amino acids 1X and 100 μg/ml kanamycin. The human 293T cell line (CRL-3216), obtained from the American Type Culture Collection (ATCC), was maintained in DMEM medium (Life Technologies) supplemented with 10% FBS, 1 mM sodium pyruvate and 1% of penicillin-streptomycin. The Raji cell line, a human B-lymphoid Epstein-Barr virus-positive cell line derived from a Burkitt’s lymphoma obtained from the AIDS Research and Reference Reagent Program (National Institute of Allergy and Infectious Disease [NIAID], National Institute of Health [NIH]), was maintained in RPMI 1640-Glutamax I medium (Life Technologies) supplemented with 10% FBS and 1% of penicillin-streptomycin. All the cells were grown at 37 °C in a humidified 95% air/5% CO2 atmosphere. Frozen PBMCs were kindly provided by Anne Van den Broeke (Unit of Animal Genomics, GIGA-R, University of Liège, Belgium) and derived from a BLV-induced B-cell leukemia (M2241L (32)).

### Plasmid constructs

The episomal plasmids pREP-luc, pREP-LTRWT-S-luc and pREP-LTR_WT_-AS-luc were previously described by our laboratory (26). Mutation of the CTCF binding site was obtained by QuickChange Site-directed Mutagenesis (Stratagene) using a pair of mutagenic oligonucleotides (**Table S2**) on a non episomal plasmid pLTR_WT_-luc (19). The resulting mutated LTR was isolated by digestion with SmaI and cloned in both orientations in the pREP-luc construct digested with BglII and blunt-ended to obtain the pREP-LTR_mCTCF_-S-luc and pREP-LTR_mCTCF_-AS-luc. Lentiviral vectors were constructed by replacement of the XbaI – AgeI fragment of a lentiviral pTRIPz plasmid (Horizon Discoveries) by the full length LTR_wt_ or LTR_mCTCF_ amplified by PCR from the pREP-LTR_wt_-S-luc or pREP-LTR_mCTCF_-S-luc constructs, respectively. The envelope pVSG-G and packaging psPAX2 vectors were obtained from the Reference Reagent Program. For all constructs, the fragments cloned were fully sequenced by Sanger sequencing.

### Chromatin Immunoprecipitation assays

ChIP assays were performed following the ChIP assay kit (EMD Millipore). Briefly, cells were cross-linked for 10 min at room temperature with 1% formaldehyde before lysis followed by sonication of their chromatin (Bioruptor Plus, Diagenode) to obtain DNA fragments of 200–400 bp. Chromatin immunoprecipitations were performed with chromatin from 6×10^6^ cells (1×10^6^ cells for the epigenetic modifications) and 5μg of antibodies (**Table S3**). Quantitative real-time PCR reactions were performed using 1/60 of the immunoprecipitated DNA and the TB Green Premix Ex Taq II (Takara). Relative quantification using standard curve method was performed for each primer pair and 96-well Optical Reaction plates were read in a StepOnePlus PCR instrument (Applied Biosystem). Fold enrichments were calculated as percentages of input values or as fold inductions relative to the values measured with IgG. Primer sequences used for quantification (**Table S2**) were designed using the software Primer 3.

### ChIP-sequencing assays

ChIP assays were performed as described above. Recovered DNA was then used for library preparation using the Ovation Ultralow System v2 kit (NuGen) following Manufacter’s instructions. Paired-end sequencing was then performed with the Illumina HiSeq 2000 instrument. More than 20 millions of single reads were obtained for all libraries. Reads were mapped to a hybrid ovine genome (OAR v3.1) containing the BLV provirus sequence (GenBank: KT122858.1) at their respective insertion site and orientation (**Table S4**) using Bowtie 2 v2.3.5.1 (43). The mapped reads with mapping quality score <15 and PCR duplicates were discarded using SAMtools v1.9 (44). CTCF peaks were called using MACS2 v2.2.5 (46) at a q-value cutoff of 0.01. CTCF motif and the orientation of each peak was identified using FIMO v5.1.1 (47) with motif MA0139.1 (48) with max-stored-scores 50000000. Bigwig files were generated with DeepTools v.3.3.2 (45) with bin length of 25 bp, extending reads to 200 bp and CPM normalization. For analysis of Rad21 and CTCF localization on the viral genome reads were mapped to a hybrid ovine genome (OAR v3.1) containing the BLV provirus sequence (GenBank: KT122858.1) at their respective insertion site and orientation with settings n1 k2 using Bowtie 2 v2.3.5.1. PCR duplicates were removed using SAMtools v1.0. Bigwig files were generated with DeepTools v.3.3.2 with bin length of 10 bp, extending reads to 200 bp and CPM normalization.

### Lentiviral production and transduction

VSV-G pseudotyped lentiviral particles were produced by transfection of 293T cells (5×10^6^ /10 cm dish), with the different LTR-containing lentiviral constructs (9 μg), the pVSV-G (2.25 μg) and the psPAX2 (6.75 μg) vectors by the calcium phosphate transfection method according to the manufacturer’s protocol (Takara). 72 hours post-transfection, viral stocks were harvested and concentrated 10x by ultracentrifugation. 293T cells (2×10^5^) were then transduced in 12well-plate with 600µl of concentrated lentiviral stock. Six hours post-transduction, medium was replaced by 1ml of complete DMEM medium. 48 hours post-transduction, puromycine selection (1µg/ml) of stably infected clones was performed for 7 days before being harvested for ChIP assays as described above.

### Transient transfection and luciferase assays

293T cells (2.5×10^5^) were co-transfected with 900ng of pREP-luc constructs and 50ng of pGL4.74 (Promega) as internal control by the calcium phosphate transfection method according to the manufacturer’s protocol (Takara). Raji cells (3×10^6^cells) were co-transfected with 600ng of pREP-luc constructs and 50ng of pGL4.74 as internal control using DEAE-dextran as described previously (23). 48 hours post-transfection, cells were lysed and luciferase activities were measured using the DualGlo-luciferase reporter assay (Promega). Results were normalized for transfection efficiency using Renilla luciferase activities and total protein concentrations.

### 4C-seq

4C-seq experiments were performed as previously described (49). Briefly, 10^7^ cells per sample were cross-linked in 2% formaldehyde for 10 min and the reaction was quenched by adding glycin at a final concentration of 0.13M. Then, cells were lysed and the harvested cross-linked chromatin was digested using the restriction enzyme DpnII (NEB). After a ligation step with the T4 DNA ligase (NEB), ligated samples were decross-linked and subjected to a second round of digestion using the restriction enzyme CviQI (NEB). Digested samples were then ligated using T4 DNA ligase and purified before being used as template for inverse PCR. Primers used for inverse PCR and library preparation are listed in **Table S2**. Products were sequenced using Illumina sequencing (Illumina NextSeq 500) and reads were mapped to a hybrid ovine genome (OAR v3.1) containing the BLV provirus sequence (GenBank: KT122858.1) at the L267 or YR2 integration sites (**Table S3**) and processed using pipe4C (49) (github.com/deLaatLab/pipe4C) with the following parameters: normalization to 1 million reads in *cis*, non-blind fragments only, window size 21, top 2 read counts removed. Coverage plots were generated using R (https://www.R-project.org/).

## RESULTS

### BLV provirus contains three relevant *in silico* CTCF binding sites

In order to assess a putative role of CTCF in the regulation of BLV gene expression and physiopathology, we first performed *in silico* analysis of the BLV proviral genome for the presence of consensus CTCF binding sites. This *in silico* analysis performed on both strand of the genome revealed the presence of several putative CTCF binding sites (**Fig. 1**). Among these putative CTCF binding sites, the two most relevant were located at the end of both BLV LTRs, spanning from nucleotide (nt) +419 to nt +514 in the 5’LTR and nt +8685 to nt +8703 in the 3’LTR, respectively (nt + 1 defined as the first nucleotide of the 5′LTR). In addition, a third CTCF site was observed in the regulatory region coding for the second exon of viral Tax and Rex (Tax/Rex E2) regulatory proteins spanning form nt +7515 to nt +7533. Of note, these putative CTCF binding sites were all located on the minus DNA strand of the BLV genome. Moreover, by performing a computational analysis of several BLV strains referenced in the NCBI database, we showed an important conservation rate of these CTCF binding sites of 81.6% (120/147) for the 5’LTR, 95.9% (141/147) for the Tax/Rex E2 and 84.3% (124/147) for the 3’LTR thereby suggesting their potentially important role in BLV life cycle and pathogenesis (**Fig. S1** and **Table S1**).

**Fig. 1:**
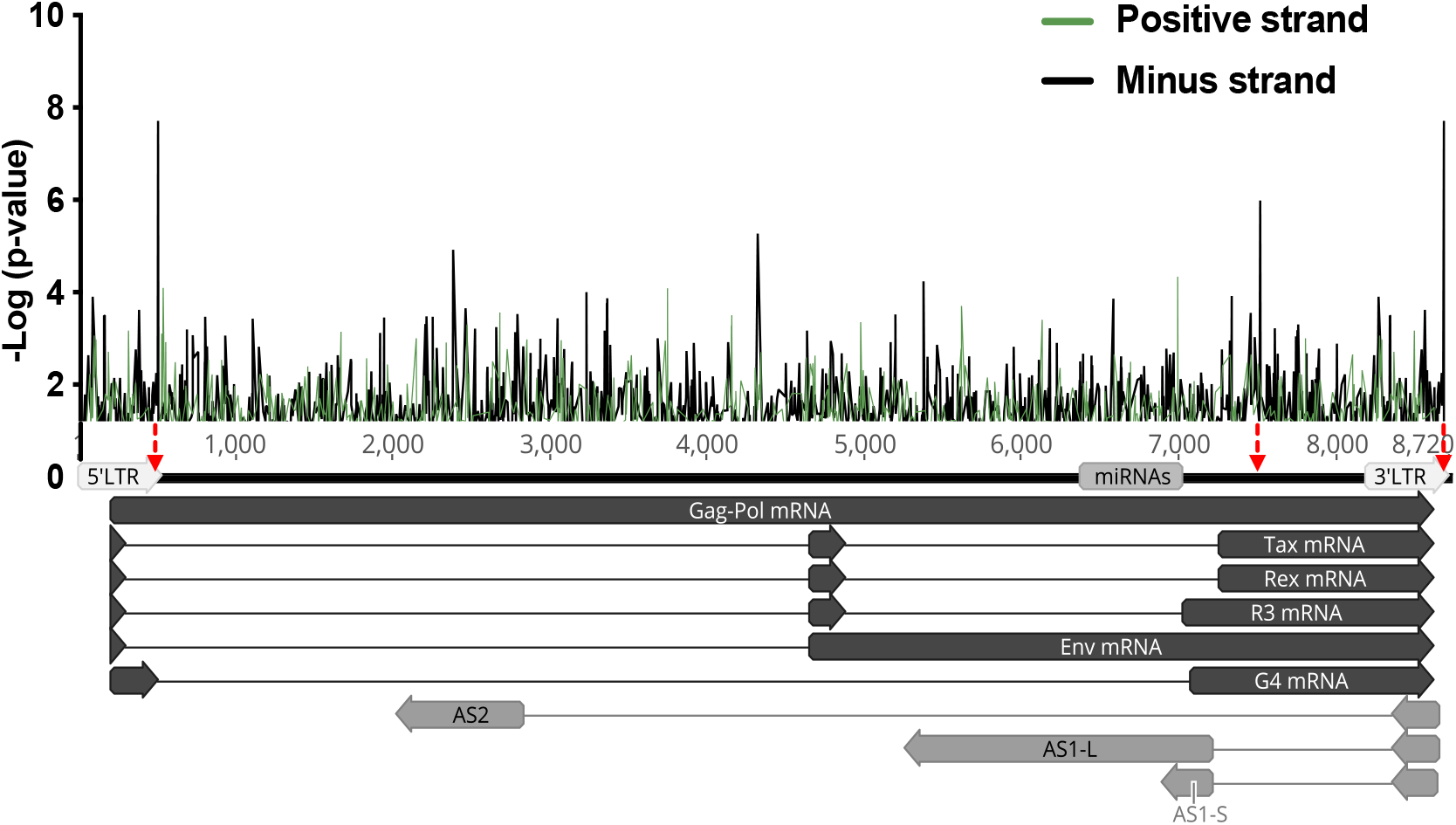
*In silico* analysis of putative CTCF binding sites along the BLV provirus. Identification of CTCF binding sites along the BLV provirus was performed using the BLV reference genome (KT122858.1) and PWMTools (74) (MA0139.1 in the MEME-derived JASPAR CORE 2018 vertebrates motif library). CTCF motifs located either on the positive or negative strand are represented by green or black lines, respectively. The three most relevant CTCF binding sites identified in silico are indicated by the red arrows.

### CTCF is recruited *in vivo* to three distinct regions along the BLV provirus

In order to determine whether the three putative CTCF binding sites identified *in silico* were able to recruit CTCF *in vivo*, we performed chromatin immunoprecipitation (ChIP) assays. To this end, we prepared chromatin from the BLV latently-infected B-lymphocytic ovine cell line L267 and immunoprecipitated CTCF using a specific antibody. As control, we used a purified IgG to measure the aspecific background. Next, purified DNA was amplified by real-time quantitative PCR (qPCR) with oligonucleotide primers hybridizing to specific regions along the BLV proviral genome and its host genomic environment (**Fig. 2**). As shown in **Fig. 2A**, we observed *in vivo* recruitment of CTCF to the end of both BLV LTRs and, to a lesser extent, to the Tax/Rex E2 region, thereby demonstrating that CTCF was recruited *in vivo* to the three regions containing the sites we identified by *in silico* analysis. To further validate the results, we next performed additional ChIP-qPCR experiments using another BLV latently-infected B-lymphocytic ovine cell line, referred to as the YR2 cell line. Of note, compared to the L267 cell line which represents an epigenetic model of viral latency, BLV infection in the YR2 cell line is latent due to two E-to K-mutations in the Tax protein, which impairs its transactivating activity (12). Our results, presented in **Fig. 2B**, showed a recruitment profile of CTCF similar to the one observed in the L267 cell line. Indeed, CTCF was also recruited to the end of both LTRs and, to a lesser extent, to the Tax/Rex E2 region. Finally, in order to validate CTCF recruitment to the BLV provirus in a more physiological model of BLV-infected cells, we performed additional ChIP-qPCR experiments using chromatin prepared from peripheral blood mononuclear cells (PBMCs) isolated from a BLV-infected sheep that developed leukemia. As shown in **Fig. 2C**, our results were in good agreement with the ones obtained in the BLV-latently infected B-lymphocytic ovine cell lines L267 and YR2, thereby confirming, in the context of BLV-latently infected primary cells, the *in vivo* recruitment of CTCF to three distinct regions along the BLV provirus: the end of both the 5’ and 3’LTRs and, to a lesser extent, to the Tax/Rex E2 region. Taken together, our results demonstrate that the 3 CTCF sites identified *in silico* are at least in part responsible for the *in vivo* recruitment of CTCF to the BLV genome, thereby providing the first evidence of a putative function of CTCF in BLV transcriptional and epigenetic regulations.

**Fig. 2:**
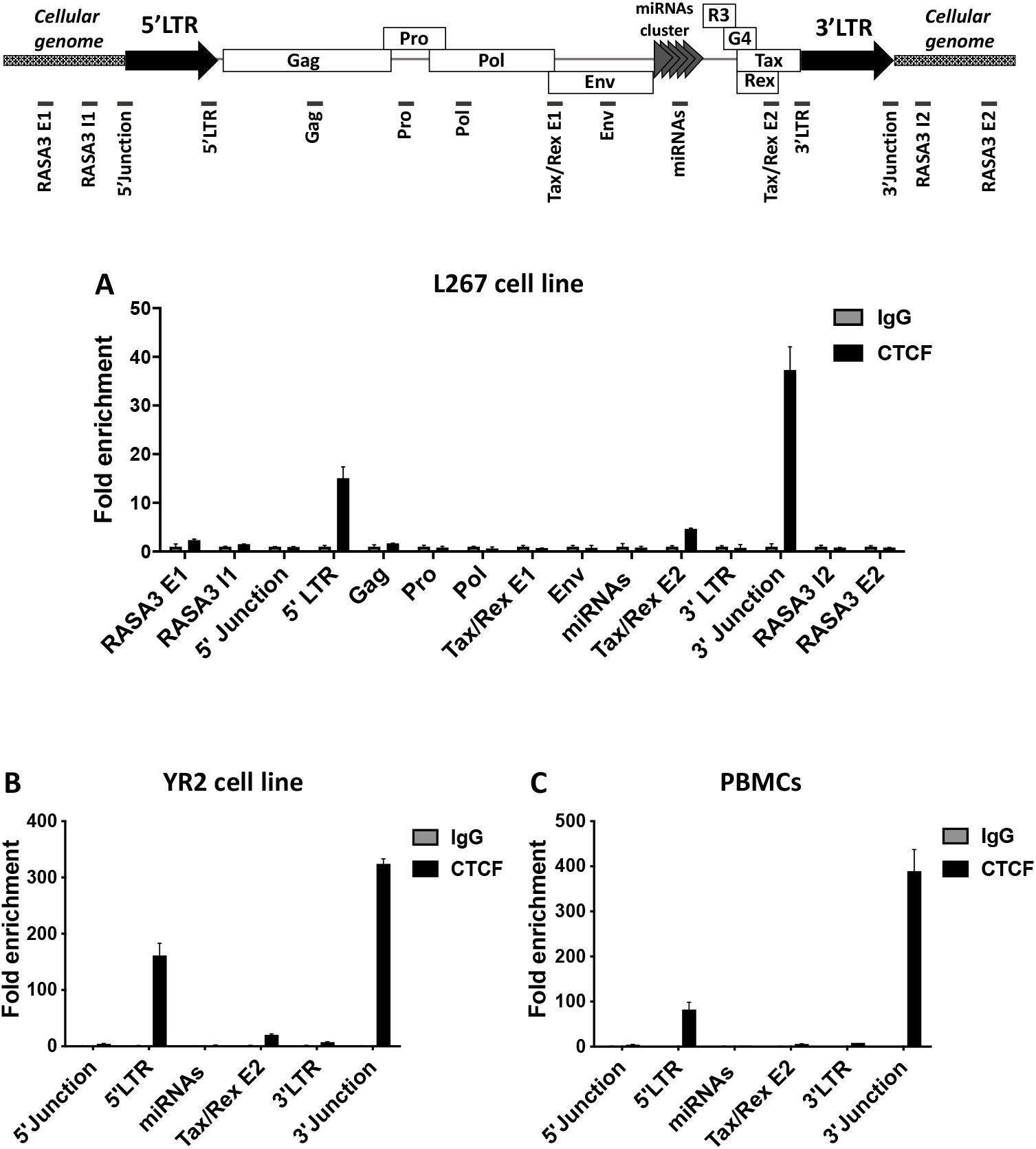
CTCF is recruited *in vivo* along the BLV provirus. Chromatin prepared from BLV-infected **(A)** L267 cell line, **(B)** YR2 cell line or **(C)** ovine PBMCs was immunoprecipitated with a specific antibody directed against CTCF or with an IgG as background measurement. Purified DNA was then amplified with oligonucleotide primers hybridizing to either the BLV genome or its host cellular surrounding DNA. Results are presented as histograms indicating fold enrichment above the value obtained with IgG, which was arbitrarily assigned the value of 1. Data are the means ± SD from one representative of at least three independent experiments.

### CTCF binding differentially regulates RNAPII-dependent transcriptional activities arising from the BLV 5’ and 3’LTRs

Previous reports have demonstrated CTCF as an important transcription factor regulating gene expression through its ability to either activate or repress gene expression, as well as to poise an enhancer region or to bring an enhancer close to its cognate promoter by formation of long-range chromatin interactions (33–36, 51). Based on the *in vivo* recruitment of CTCF to the 5’ and 3’ LTRs, which respectively exhibit sense and antisense RNAPII-dependent promoter activities, we decided to investigate the putative functional role played by CTCF in these two transcriptional activities. To this end, we designed point mutations in the CTCF binding motif of BLV LTR carefully avoiding the insertion of mutations in *cis*-regulatory elements previously described as critical for either the sense or antisense promoter activities (18–26) (**Fig. 3A**). We then validated the effects of the designed mutations on CTCF recruitment *in vivo*, by performing ChIP assays after transduction of either the wild-type or the CTCF-mutated BLV LTR. Briefly, lentiviral particles containing either the wild-type or the CTCF-mutated LTR were produced and used to transduce HEK293T cell line. Transduced cells were selected using puromycin and assayed for ChIP experiments. Prepared chromatin was immunoprecipitated using a specific antibody directed against CTCF or with purified IgG to measure aspecific background. Next, purified DNA was amplified by specific oligonucleotide primers surrounding the CTCF binding site of the LTRs. As positive and negative controls, we also designed specific oligonucleotide primers hybridizing to two cellular genomic regions, MDM2 and GAPDH, previously reported to either recruit or not CTCF (52). As shown in **Fig. 3B**, the designed mutations completely abolished the *in vivo* recruitment of CTCF to the BLV LTR binding site. In order to evaluate the functional role of CTCF on both 5’LTR sense or 3’LTR antisense RNAPII-dependent transcriptional activities, we cloned either the wild-type or CTCF-mutated BLV LTR upstream of the *Firefly* luciferase (luc) reporter gene in both the 5′ and 3′ orientations (mimicking the sense and antisense RNAPII-dependent transcriptional activities, respectively) into a modified pREP10 episomal vector (pREP-luc). Of note, using episomally replicating luciferase reporter constructs has the advantage to display the hallmarks of proper chromatin structure when transiently transfected into cells (53, 54), an important feature allowing the study of transcriptional effects while taking into account the chromatin structure. The reporter constructs (referred to as pREP-LTR_wt_ S-luc, pREP-LTR_wt_ AS-luc, pREP-LTR_mCTCF_ S-luc and pREP-LTR_mCTCF_ AS-luc) were then transiently transfected into the HEK293T cell line. Forty-hours post-transfection, cells were lysed and assayed for luciferase activity. As shown in **Fig. 3C**, the RNAPII-dependent sense transcription from the LTR promoter was significantly increased by 5.45-fold (p-value = 0.0005) when CTCF recruitment is abolished, thereby suggesting a repressive effect of CTCF on the 5’LTR promoter activity. However, when cloned in the antisense orientation, the CTCF-mutated LTR promoter activity is significantly decreased by 4-fold (p-value = 0.0005), thereby suggesting an activating effect of CTCF on the 3’LTR promoter activity. Thus, these results showed a differential impact of CTCF binding to the BLV LTR depending on its orientation. In addition, to confirm our results in a more relevant context of natural target cells corresponding to B cells, we performed additional transient transfection experiments into the human B-lymphoïd cell line Raji, using the same experimental procedure (**Fig. 3D**). Similar to what we observed in 293T cells, our results showed a differential functional role of CTCF, favoring the RNAPII-dependent antisense transcription while repressing the RNAPII-dependent sense transcription **(Fig. 3D)**. Taken together, our results demonstrate an important functional role of CTCF in the BLV LTR-driven promoter activities in the context of transient transfection of episomal constructs.

**Fig. 3:**
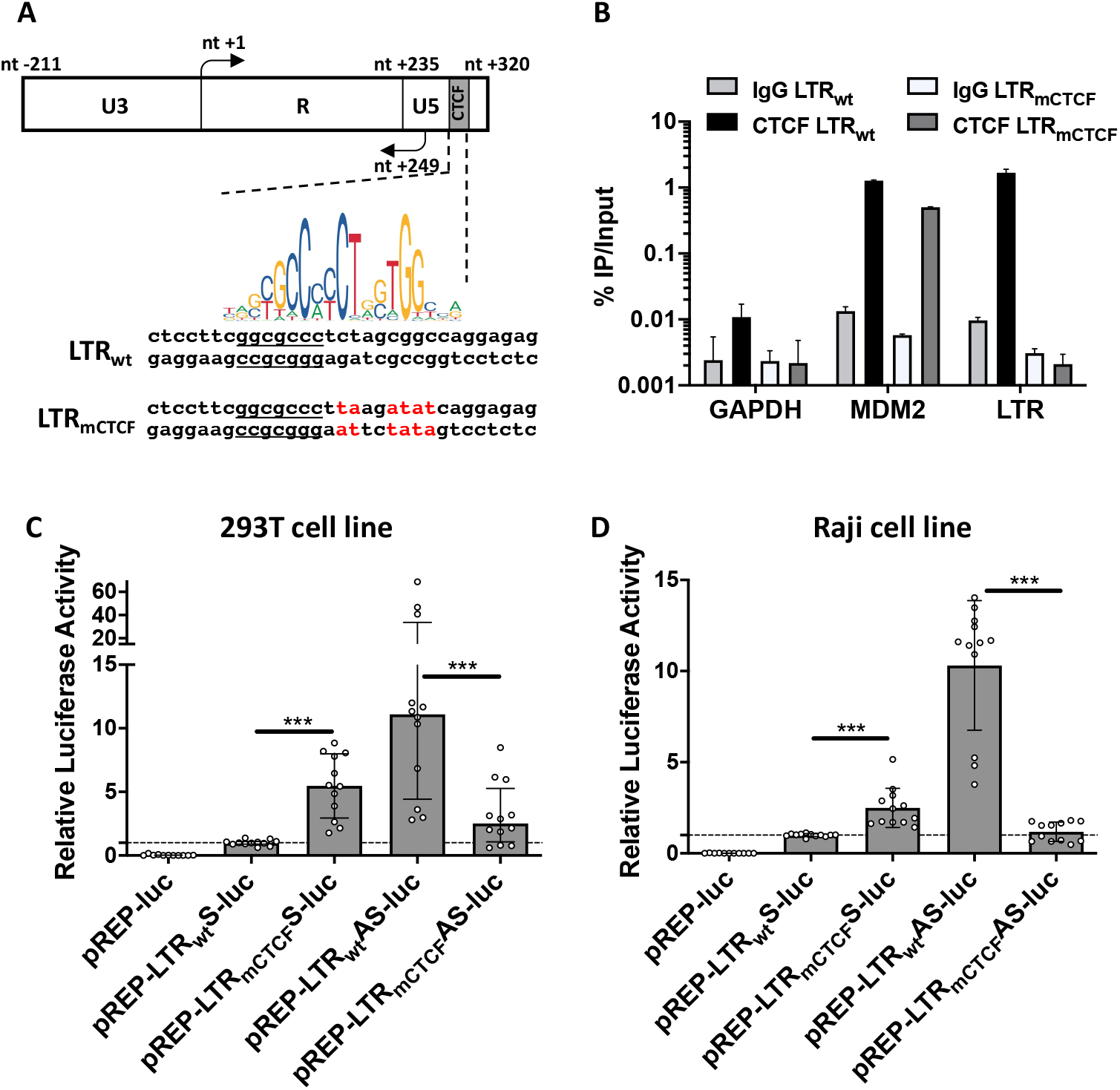
CTCF binding differentially impacts BLV LTR promoter activities. **(A)** Schematic representation of the BLV LTR with the identified TSS for either the sense (nt +1) or antisense (nt +249) transcription. CTCF logo is indicated below as well as the wild-type LTR (LTR_wt_) or mutated LTR (LTR_mCTCF_) containing mutations abolishing CTCF binding. The closest *cis-*regulatory element (BRE) is underlined (26). **(B)** Chromatin prepared from 293T cells stably transduced with either the wild-type BLV LTR or the mutated LTR was immunoprecipitated with a specific antibody directed against CTCF or with an IgG as background measurement. Purified DNA was then amplified with oligonucleotide primers hybridizing to either the BLV LTR or host cellular genes (GAPDH and MDM2). Results are presented as histograms indicating percentages of immunoprecipitated DNA compared to the input DNA (% IP/Input). Data are the mean ± SD from one representative of at least three independent experiments. **(C)** 293T or **(D)** Raji cells were transiently co-transfected with 600 ng of pREP-luc, pREP-LTR_wt_-S-luc, pREP-LTR_mut_-S-luc, pREP-LTR_wt_-AS-luc or pREP-LTRmut-AS-luc constructs and 50 ng of pRL-TK. Forty-eight hours after transfection, cells were collected, lysed and both *Firefly* and *Renilla* luciferase activities were measured. Results are presented as histograms indicating relative luciferase activities (RLU) compared to the value obtained with the pREP-LTR_wt_-S-luc construct which was assigned to the value of 1. Data are the medians ± interquartile of at least three independent experiments. The Wilcoxon signed-rank (value of 1) or the Mann-Whitney statistical tests were used for sense and anti-sense orientations, respectively, with P > 0.05 = ns ; P ≤ 0.05 = * ; P ≤ 0.01 = ** ; P ≤ 0.001 = ***

### CTCF determines a specific epigenetic profile along BLV proviral genome

CTCF exhibits a broad range of functions and is enriched at boundaries of topologically associated domains (TADs). These TADs are regions where DNA contacts preferentially occur and which are often characterized by a similar epigenetic landscape (36, 53). Even though the direct functional role of CTCF in delimitating epigenetic borders is still under debate (55), several studies have linked CTCF to the maintenance of distinct epigenetic profiles (56–58).

We have previously reported the epigenetic profile along the three BLV promoters in the context of the BLV latently-infected B-lymphocytic ovine cell line L267 (26). Here, to investigate whether CTCF could contribute to the establishment of a putative epigenetic border along the entire BLV provirus, we performed ChIP-qPCR experiments using chromatin from the BLV latently-infected B-lymphocytic ovine cell line L267 which was immunoprecipitated with specific antibodies directed against several histone post-translational modifications. As control, we used a purified IgG to measure the aspecific background. Purified DNA was amplified by qPCR with oligonucleotide primers hybridizing to specific regions along the BLV provirus and its host genomic environment. As shown in **Fig. 4A**, we observed an enrichment of histone post-translational modification marks associated with active transcription i.e. acetylated histones (AcH3), acetylated histone 3 lysine 9 (H3K9ac), acetylated histone 3 lysine 27 (H3K27ac), di-methylated histone 3 lysine 4 (H3K4me2), tri-methylated histone 3 lysine 4 (H3K4me3). These epigenetic marks were encompassing the proviral region delimitated by CTCF binding to the Tax/Rex E2 and 3’LTR regions. Of note, this positive epigenetic signature correlated with the constitutive antisense RNAPII-dependent transcription arising from the 3’LTR (26, 29). Moreover, in agreement with transcriptional silencing affecting the BLV 5’LTR, these results showed an absence of positive epigenetic marks at the 5’LTR. Overall, our results demonstrated an epigenetic signature clustered by the regions to which CTCF is recruited.

**Fig. 4:**
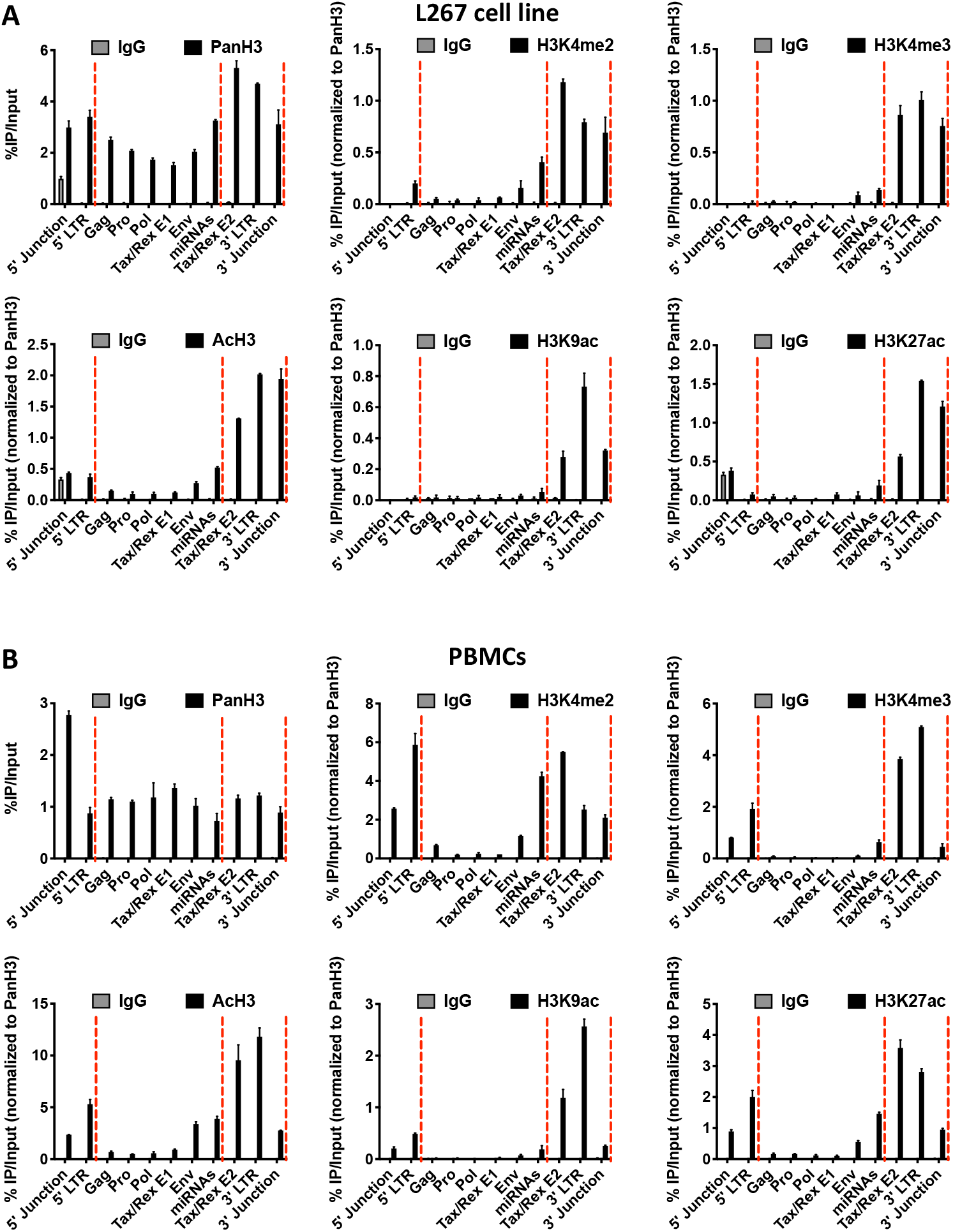
CTCF defines a specific epigenetic profile along the BLV proviral genome. Chromatin prepared from BLV infected **(A)** L267 cells or **(B)** ovine PBMCs was immunoprecipitated with specific antibodies directed against histone H3 (PanH3), different histone post-translational modifications (H3Kme2, H3Kme3, AcH3, H3K9ac, H3K27ac, H3k27me3, H3K36me3) or with an IgG as background measurement. Purified DNA was then amplified with oligonucleotide primers hybridizing to the BLV proviral genome. Results are presented as histograms indicating percentages of immunoprecipitated DNA compared to the input DNA (% IP/Input) normalized to PanH3. Data are the means ± SD from one representative of at least three independent experiments.

Next, in order to strengthen our results showing a potential role of CTCF in delimiting an epigenetic border along the BLV provirus, we performed additional ChIP-qPCR experiments in the context of another BLV latently-infected B-lymphocytic cell line YR2 (**Fig. S2**) and of a more physiological context of BLV infection corresponding to BLV-infected ovine PBMCs (**Fig. 4B**). Our results confirmed an enrichment of positive histone post-translational modifications which was clustered by the two CTCF binding sites located in the Tax/Rex E2 region and in the 3’LTR, respectively. However, regarding the epigenetic profile along the 5’LTR and in opposition to what we observed in the L267 cell line, an accumulation of positive histone marks was observed in the 5’LTR in both the YR2 cell line and in PBMCs. These observations were in agreement with previous results describing a weaker latency state compared to the L267 cell line associated with a low basal level of RNAPII-dependent sense transcription (13, 26, 32). However, since these positive epigenetic marks were not spreading downstream of CTCF binding to the 5’LTR, CTCF could also establish an epigenetic boundary contributing to viral latency, a situation in agreement with our data obtained by transient transfection assays (**Fig. 3C** and **3D**).

Taken together, our results provide evidence that CTCF might be implicated in the maintenance of an active RNAPII-dependent antisense transcription, by the delimitation of an active epigenetic landscape, while maintaining the RNAPII-dependent 5’LTR promoter activity silent, by preventing the spread of positive epigenetic marks.

### The BLV provirus induces abnormal chromatin loops with its host cellular environment

One of the most studied role of CTCF is the organization of 3D chromatin architecture, notably resulting in long-range interactions, forming structural domains and bringing enhancers/silencers close to their cognate promoters by forming chromatin loops between two CTCF binding sites (33, 34, 36, 59, 60). The formation of these chromatin loops requires the co-recruitment of the cohesin multiprotein complex to the CTCF binding sites mediated by a mechanism called loop extrusion (61). In this context, we decided to assess whether the BLV provirus was able to induce the formation of abnormal chromatin loops with the host genomic environment. To this end, we first studied the co-recruitment of the cohesin complex to the previously identified regions of CTCF recruitment binding sites by performing ChIP-qPCR experiments. Chromatins prepared from the two BLV latently-infected B-lymphocytic cell lines L267 and YR2 or from BLV-infected ovine PBMCs were immunoprecipitated with a specific antibody targeting the Rad21 subunit of the cohesin complex or with a purified IgG to measure the aspecific background. Purified DNA was next amplified by qPCR using oligonucleotide primers hybridizing to specific regions along the BLV genome (**Fig. 5**). In all the cellular models of BLV infection tested, we observed the *in vivo* recruitment of Rad21 to the end of both the 5’ and 3’LTRs (**Fig. 5A-C**), thereby suggesting a cooperative role of CTCF and cohesin in the formation of chromatin loops involving the BLV CTCF binding regions.

**Fig. 5:**
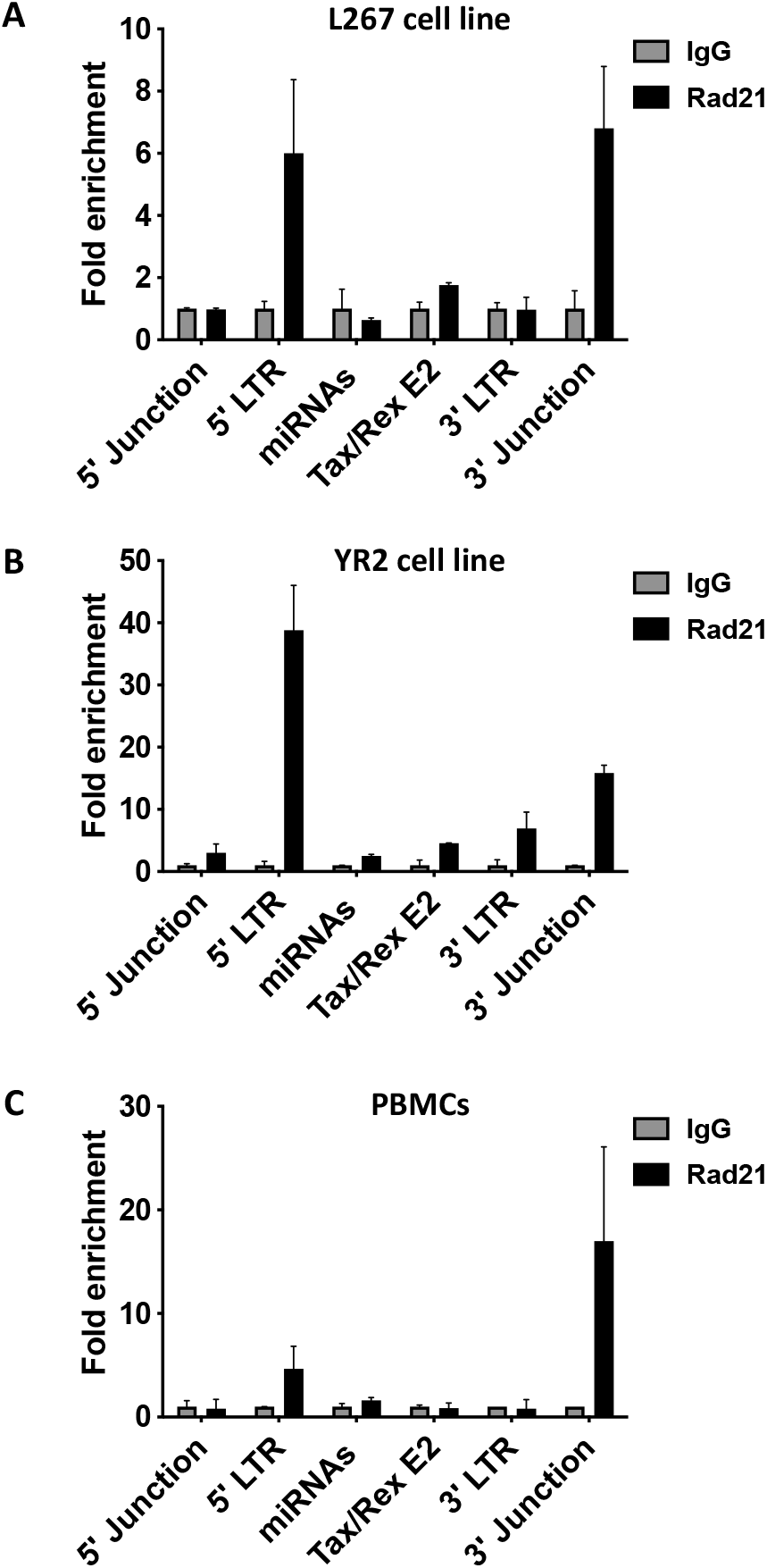
The Rad21 subunit of the cohesin multiprotein complex is co-recruited to the BLV CTCF binding sites. Chromatin prepared from BLV infected **(A)** L267 cells, **(B)** YR2 cells or **(C)** ovine PBMCs was immunoprecipitated with specific antibodies directed against Rad21 or with an IgG as background measurement. Purified DNA was then amplified with oligonucleotide primers hybridizing to either the BLV proviral genome or its host cellular surrounding DNA. Results are presented as histograms indicating fold enrichment above the value obtained with IgG, which was arbitrarily assigned the value of 1. Data are the means ± SD from one representative of at least three independent experiments.

In order to assess whether the integrated BLV provirus was able to establish 3D chromatin structures with its host genome, we performed circular chromosome conformation capture (4C) experiments followed by high-throughput sequencing (4C-seq), a technique allowing the identification of physical contacts between a specific DNA region, called the viewpoint, and the entire cellular genome. Using the BLV-latently infected B-lymphotropic ovine cell lines L267 and YR2, we designed three specific viewpoints encompassing each BLV CTCF binding sites. As shown in **Fig. 6**, we clearly identified the presence of chromatin loops between the BLV provirus and its host genome in both the L267 and YR2 cell lines using a viewpoint encompassing the CTCF binding site located in the 3’LTR. In the context of the L267 cell line, we observed a strong interaction between the BLV provirus and at least one genomic region located approximatively 60kb downstream of the BLV integration site (IS) (**Fig. 6A**). To study the possible role of CTCF and cohesin in the formation of these chromatin loops, we performed ChIP-seq experiments to assess the genome-wide occupation of CTCF and cohesin. Among other regions, we observed that CTCF and cohesin co-localize at the genomic region physically interacting with the BLV provirus (**Fig. 6A**). Similar 4C-seq profiles were obtained when we used the two other viewpoints encompassing the CTCF binding sites located in the 5’LTR and in the Tax/Rex E2 region, respectively (**Fig. S3**), thereby demonstrating that the proviral regions induced local topological changes in the host genome. In the context of the YR2 cell line, having the viral DNA integrated at another location in the genome, the 4C-seq data again showed multiple physical contacts between the viral CTCF sites and surrounding CTCF sites present in the host genome **(Fig. 6B; Fig. S3**). Altogether, our results demonstrated that the integrated BLV provirus induces ectopic chromatin loops with its host cellular environment. We speculate that this provides a new mechanism used by this leukemogenic retrovirus to perturbate the host 3D cellular genome organization and, as a consequence, potentially the host cellular transcriptome.

**Fig. 6:**
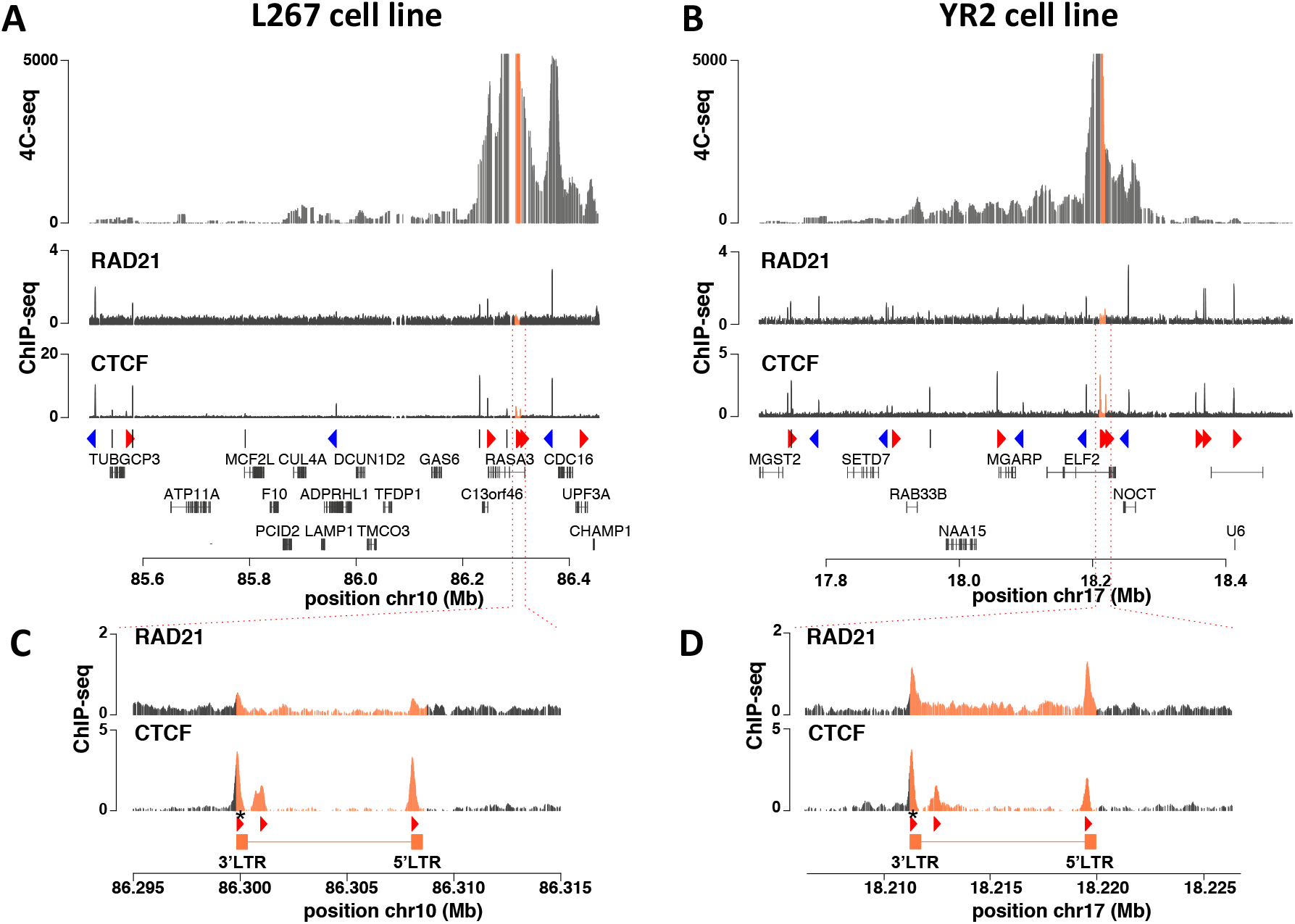
Chromatin contacts are established between BLV and cellular genomic regions. 4C-seq contact profile (average of 2 biological replicates) in the context of **(A)** L267 cells or **(B)** YR2 cells using BLV CTCF binding site of the 3’LTR as viewpoint. Below the 4C plots are shown the Rad21 and CTCF ChIP-seq profiles. CTCF binding motifs orientation is indicated by red (forward) or blue (reverse) arrows or by gray bars (undetermined) as well as the position of host cellular genes, with respect to the previously identified BLV integration site. Zoom-in of the Rad21 and CTCF ChIP-seq profiles around BLV integration site are shown for **(C)** L267 cells or **(D)** YR2 cells, as well as the provirus orientation and viewpoint localization (black asterisks).

## DISCUSSION

Despite the well-described BLV latency affecting the BLV 5’LTR promoter activity and preventing the viral genes expression, it has been demonstrated that the transcriptional network regulating BLV gene expression is more complex than initially assumed since two additional promoter activities have been discovered with an RNAPIII-dependent promoter activity, responsible for an active transcription of 10 viral micro-RNAs, and an RNAPII-dependent antisense transcription, arising from the 3’LTR and allowing high expression levels of non-coding antisense transcripts (26–29). Altogether, these findings have highlighted the deep complexity of BLV transcriptional regulation and paved the way to discover new mechanisms allowing the tumoral development. In the present report, we studied the implication of the cellular protein CTCF in this complex transcriptional network as well as its role in the establishment of a specific epigenetic profile along the BLV provirus and in the perturbation of the 3D chromatin organization of infected cells. First, we identified *in silico* three highly conserved CTCF binding sites along the BLV provirus and demonstrated the *in vivo* recruitment of CTCF in BLV latently-infected B-lymphocytic ovine cell lines as well as in BLV-infected primary cells isolated from a leukemic sheep (**Fig. 1** and **2**). Then, we investigated whether CTCF could act as a transcriptional regulator of either the sense or the antisense RNAPII-dependent transcriptional activities by introducing point mutations in the BLV LTR CTCF binding sites, thereby preventing its *in vivo* recruitment (**Fig. 3A** and **3B**). Importantly, we demonstrated that CTCF exhibits opposite effects depending on the orientation of the RNAPII-dependent transcription. Indeed, our results showed that CTCF binding to the 5’LTR inhibits the RNAPII-dependent sense transcription, while CTCF binding to the 3’LTR enables the maintenance of an active RNAPII-dependent antisense transcription (**Fig. 3**). Of note, our data showing that CTCF is an activator of the 3’LTR promoter activity are in good agreement with the literature reporting that CTCF acts as a transcriptional activator when bound upstream of the transcription start site (TSS) in a concordant orientation with the transcribed gene (55, 62), as observed here in the context of the BLV 3’LTR **(Fig. 3A)**, where CTCF is located approximatively 45bp upstream of the TSS responsible for the antisense transcription (26, 29). Regarding the role of CTCF recruitment to the 5’LTR, our results demonstrated a repressive effect on the RNAPII-dependent sense transcriptional activity, thereby indicating that CTCF could contribute to the establishment of BLV latency, a major feature to escape the host immune system while allowing tumoral development. Overall, the present study demonstrates a critical role of CTCF in the regulation of BLV transcriptional network by maintaining the 5’LTR promoter latent, while allowing expression of antisense transcripts arising from the 3’LTR.

Among the multiple roles of CTCF described in the literature, its recruitment to epigenetic boundaries as well as its implication in the delimitation of topologically associated domains has been widely studied. In the retrovirology field, a previous study has reported a single CTCF binding site in the HLTV-1 pX regulatory region, acting as an epigenetic border favoring antisense transcription through the maintenance of a positive epigenetic landscape (38). However these latter results were then refuted by a more recent publication (39). Here, we studied the putative role of CTCF in the establishment of a specific epigenetic signature along the BLV provirus. Our results clearly showed a distinct epigenetic profile with positive epigenetic marks spreading from the 3’LTR to the pX region, while these marks strongly decreased from this latter region to the 5’LTR. In addition, in the context of BLV-infected PBMCs, we also showed that these positive epigenetic marks were enriched along the 5’LTR but immediately dropped downstream. Interestingly, by comparing the epigenetic profiling along the BLV provirus with our data demonstrating the *in vivo* recruitment of CTCF, we hypothesized that CTCF recruitment to the 5’LTR could prevent the spread of positive epigenetic marks, allowing the reinforcement of BLV latency while in the meantime favoring the antisense transcription through the establishment of a positive epigenetic landscape spreading from its binding site in the pX region to the 3’LTR. Altogether, our results demonstrate that CTCF could contribute to the BLV 5’LTR silencing by acting not only at the transcriptional level but also at the epigenetic level. In contrast, CTCF could favor the RNAPII-dependent antisense transcription by acting as a transcriptional activator and a positive epigenetic regulator, thereby identifying CTCF as a new key regulator of BLV gene expression.

Despite the latest discoveries about the putative role of either BLV miRNAs (30, 31) or chimeric viral-host transcripts arising from the 3’LTR (32), the precise mechanisms underlying BLV-induced leukemogenesis remain incompletely understood and could be a multifactorial phenomenon resulting from a combination of multiple cellular deregulations, thereby leading to the irreversible cellular transformation of infected B cells. Since CTCF has also been demonstrated to contribute to the chromatin organization at the 3D level, we investigated whether such chromatin loops, induced by the inserted BLV CTCF binding sites, could contribute to BLV-associated pathogenesis. After performing 4C-seq experiments in the two BLV latently-infected B-lymphocytic ovine cell lines L267 and YR2, we showed that BLV was able to contact its host cellular environment, through the formation of viral-host chromatin loops established after the co-recruitment of CTCF and of the cohesin multiprotein complex to the LTRs, as demonstrated by our ChIP-qPCR and ChIP-seq data (**Fig. 2, Fig. 5** and **Fig. 6**). Together, these results indicate that the viral CTCF binding sites, randomly inserted into the host cellular genome following BLV integration, might modify the natural host 3D chromatin architecture and, as consequence, deregulate the expression of cellular genes that are critical for cell homeostasis and favor tumoral development. Indeed, deregulation of cellular gene expression through disruption of chromatin loops involving CTCF has frequently been associated with several diseases, including cancers (63–67). Moreover, the LTR is a well-described enhancer sequence (18–25) and therefore, bringing the LTR enhancer activity close to a cellular gene through such viral-host chromatin loops may directly deregulate gene expression, a situation already observed in the context of KSHV (68, 69), EBV (70) and where viral CTCF-mediated chromatin loops determine viral promoter usage and latency type. Our 4C-seq data in L267 ovine cells revealed a major contact between the BLV provirus and the host genome identified in the vicinity of the CDC16 gene, which codes for a member of the anaphase-promoting complex (APC/C) known for its involvement in cell cycle progression through control of progression of mitosis (71). By comparison, in the YR2 cell line, we observed a major contact made with the NOCT gene, encoding for a phosphatase involved in the regulation of cellular metabolism (72). Regarding the other minor contacts, they encompass other genes such as MGARP, NAA15, RAB338 and SETD7. Interestingly, SETD7 codes for an histone methyltransferase which acts as a tumor suppressor and was demonstrated to be downregulated in lung cancer (73). Altogether, we highlighted several putative deregulated cellular targets which will be further investigated to evaluate their role in BLV-associated physiopathology. Of note, since BLV integration is a random process, the formation of such viral-host chromatin loops will also be a random process, a key feature which could explain that BLV-associated leukemogenesis occurs in only 5% of infected animals. Associated with the previously described roles of BLV miRNAs and chimeric viral-host transcripts in host transcriptome deregulation, we identify in the present report a potential critical step leading to tumor development.

Finally, a major feature of BLV-induced B-cell tumors is the presence of an integrated provirus containing large deletions, often affecting the 5’LTR and therefore also explaining the viral latency phenomenon. Interestingly, a recent publication from the group of Anne Van den Broeke has demonstrated that the 3’LTR, in which we observed the highest co-recruitment of CTCF and Rad21, is always conserved in tumor isolates, thereby demonstrating the critical importance of this proviral region in BLV physiopathology (32). Moreover, it has been observed that cancer driver genes are enriched upstream of BLV integration sites and are perturbated by the RNAPII-dependent antisense transcription arising from the 3’LTR (32). While these genes are relatively closer to the integrated provirus, the formation of viral-host chromatin loops, highlighted in the present report, could be an additional mechanism used by BLV to perturbate its host transcriptome, but at long-range distances. This hypothesis is strengthened by the fact that CTCF binding sites along the BLV provirus are in convergent orientation with the sites found upstream of the BLV integration site, thereby rendering the formation of chromatin loops highly probable (60).

Taken together, the present report identifies CTCF as a new regulator of BLV gene expression and associated physiopathology, by acting both at the transcriptional and epigenetic levels to specifically allow the RNAPII-dependent antisense transcription, while maintaining the 5’LTR promoter latent. In addition, our results pave the way for a critical role of CTCF binding along the BLV provirus to induce viral-host chimeric chromatin loops with, as consequence, the deregulation of the host cellular transcriptome and the trigger of tumoral development. Therefore, our results provide new insights into BLV transcriptional and epigenetic regulation as well as into BLV-associated leukemogenesis, thereby highlighting mechanisms used by retroviruses to regulate their own expression and to alter the host cellular transcriptome. In addition, a better understanding of these mechanisms will allow us to identify new strategies to cure BLV infection and, to a larger extent, decrease the economical losses in endemic countries.

## DATA AVAILABILITY

All data are available in the main text, in the supplementary materials or upon request. ChIP-seq and 4C-seq data have been deposited in the Gene Expression Omnibus with accession codes GSE178482 and GSE178483, respectively.

## SUPPLEMENTARY DATA

Supplementary Data are available at NAR online.

## ACKNOWLEDGEMENT

We acknowledge A. Van den Broeke (GIGA, University of Liège, Belgium) for generously providing BLV-infected ovine primary cells used in this work. We acknowledge the Lausanne Genomic Technology Facility of the University of Lausanne for performing high-throughput sequencing experiments and bioinformatical analyses. We thank all the members of the Van Lint laboratory for discussions and critical reading of the manuscript.

