## Supplementary file for "Implication of the cellular factor CTCF in the regulation of Bovine Leukemia Virus latency and tridimensional chromatin organization"

**This PDF file includes:**

Figs. S1 to S3
Tables S1 to S4

5’LTR

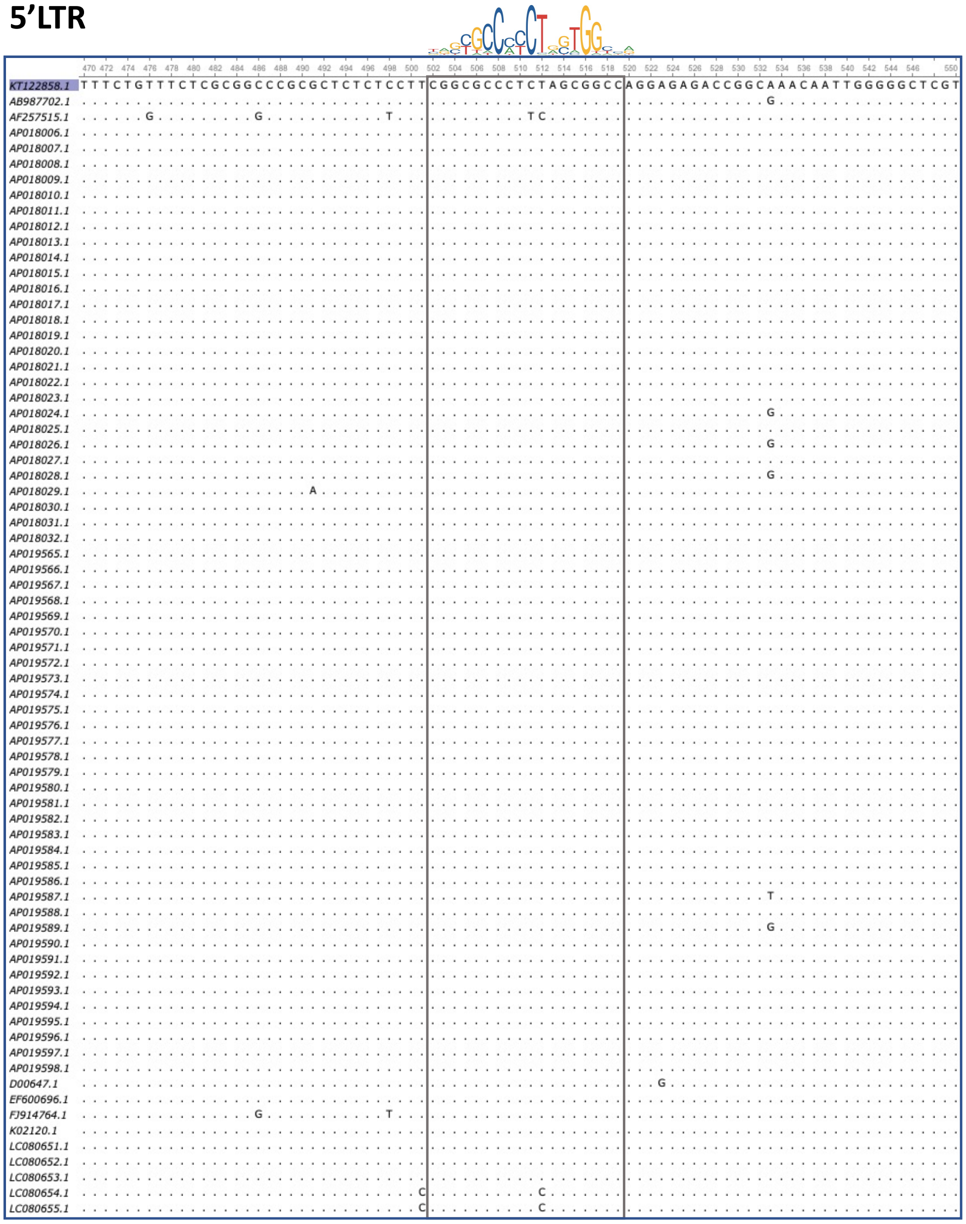


**Fig. S1 : Conservation of the CTCF DNA motif in the BLV genome.** 147 full length BLV genomes publicly available on NCBI were aligned using clustalo (1.2.4). Alignments were visualized using Unipro Ugene. CTCF consensus binding motif MA0139.1 is shown.

5’LTR

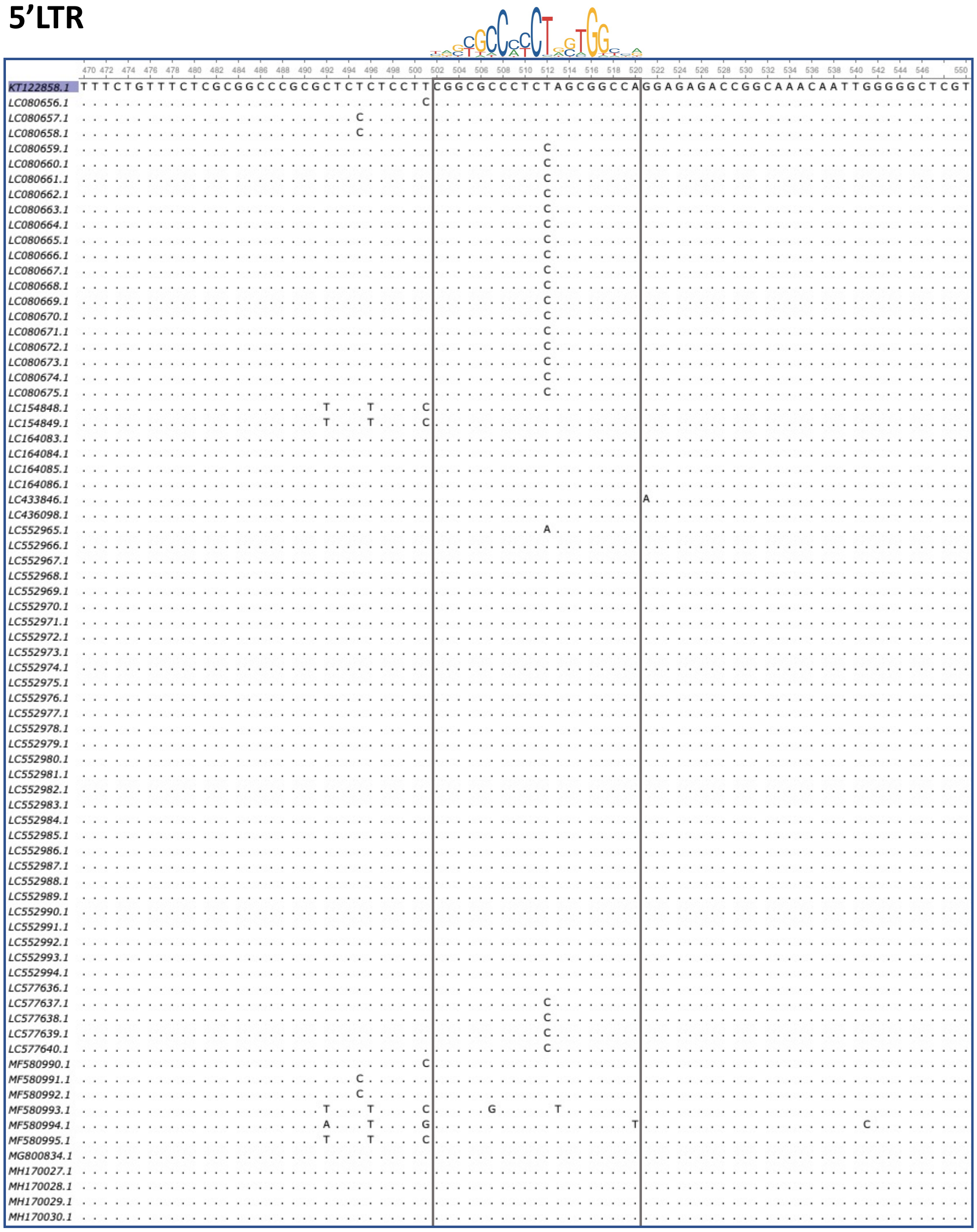

**Fig. S1 (continued).**

### Tax/Rex E2

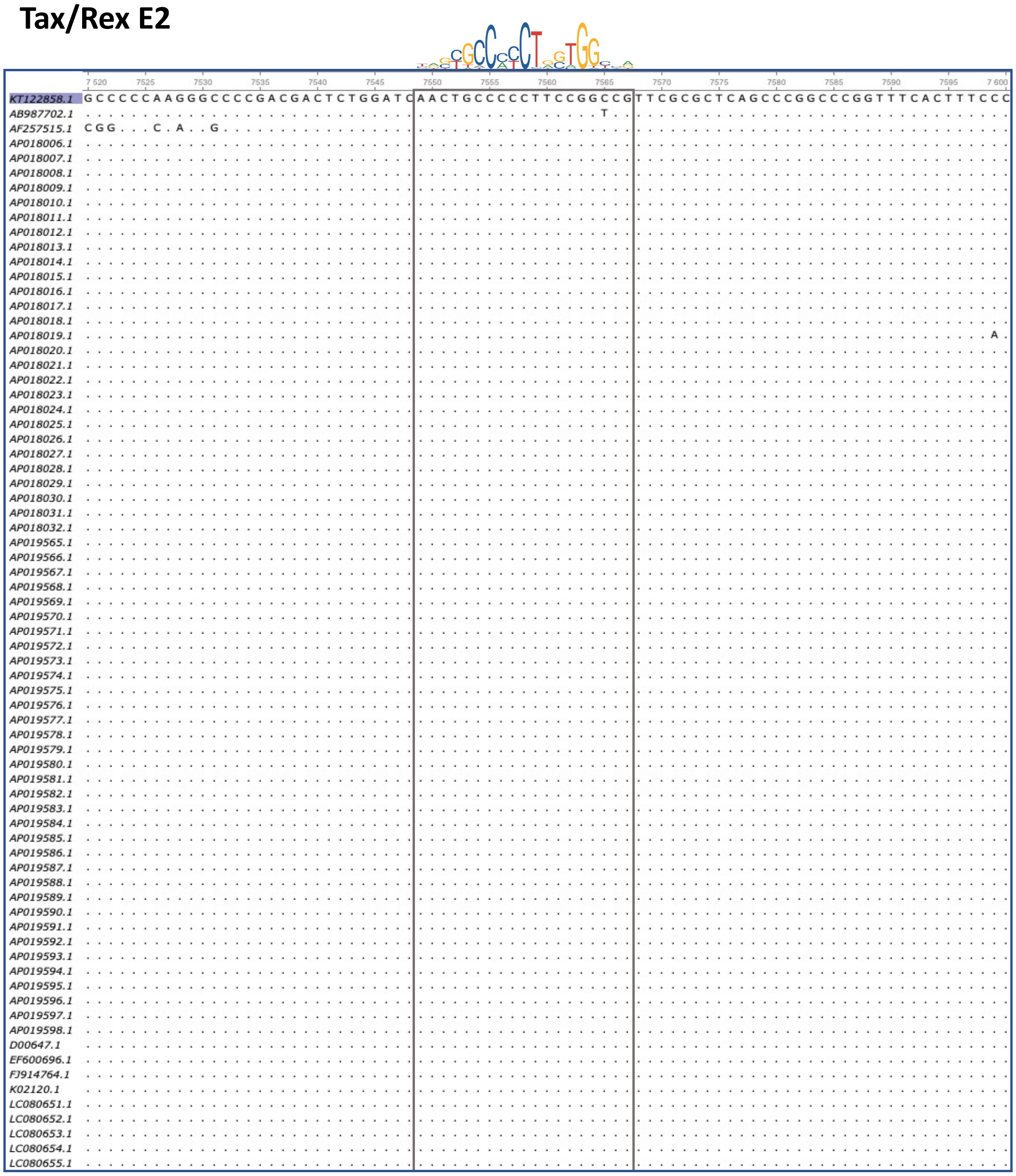

**Fig. S1 (continued).**

### Tax/Rex E2

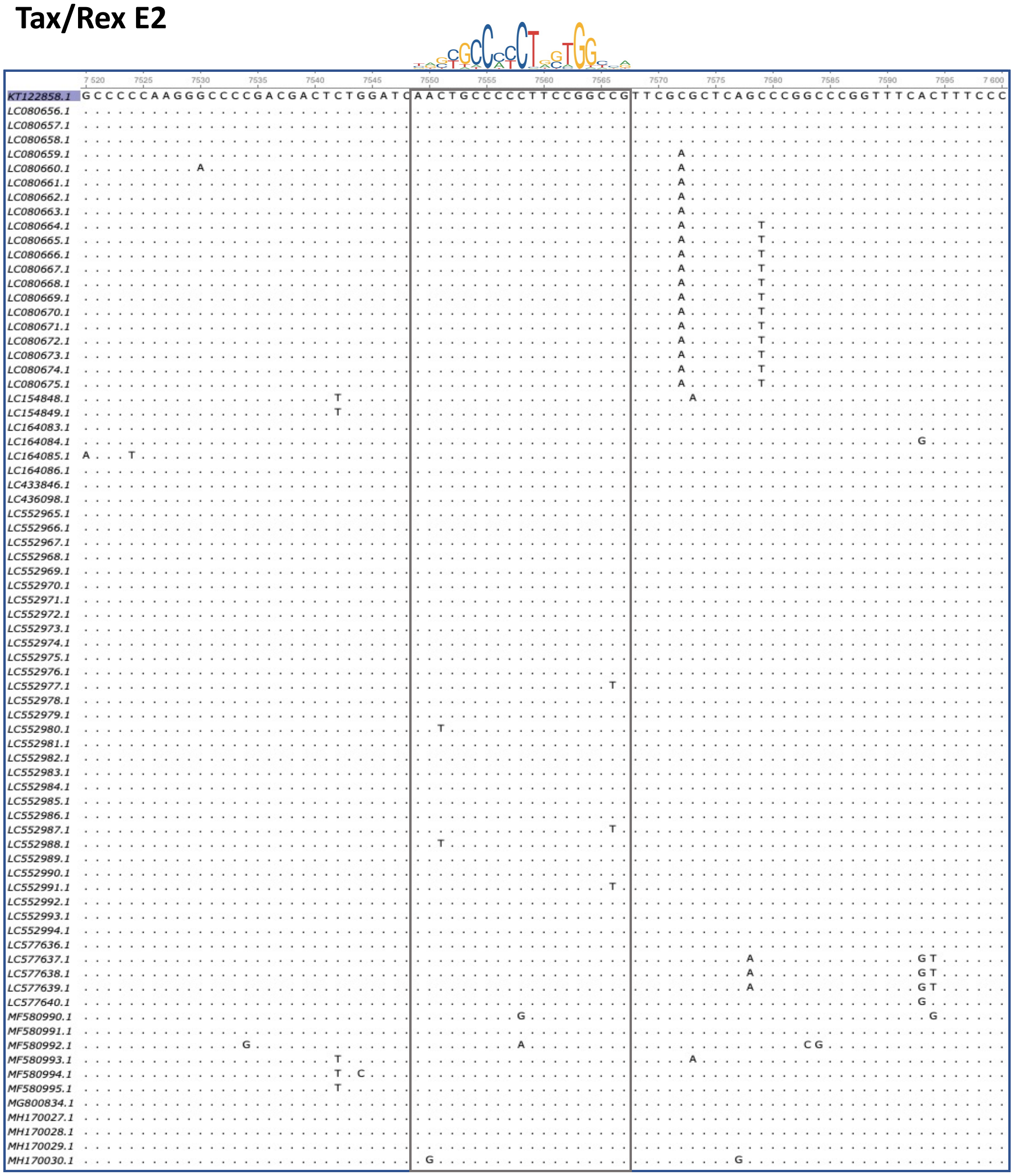

**Fig. S1 (continued).**

### 3’ LTR

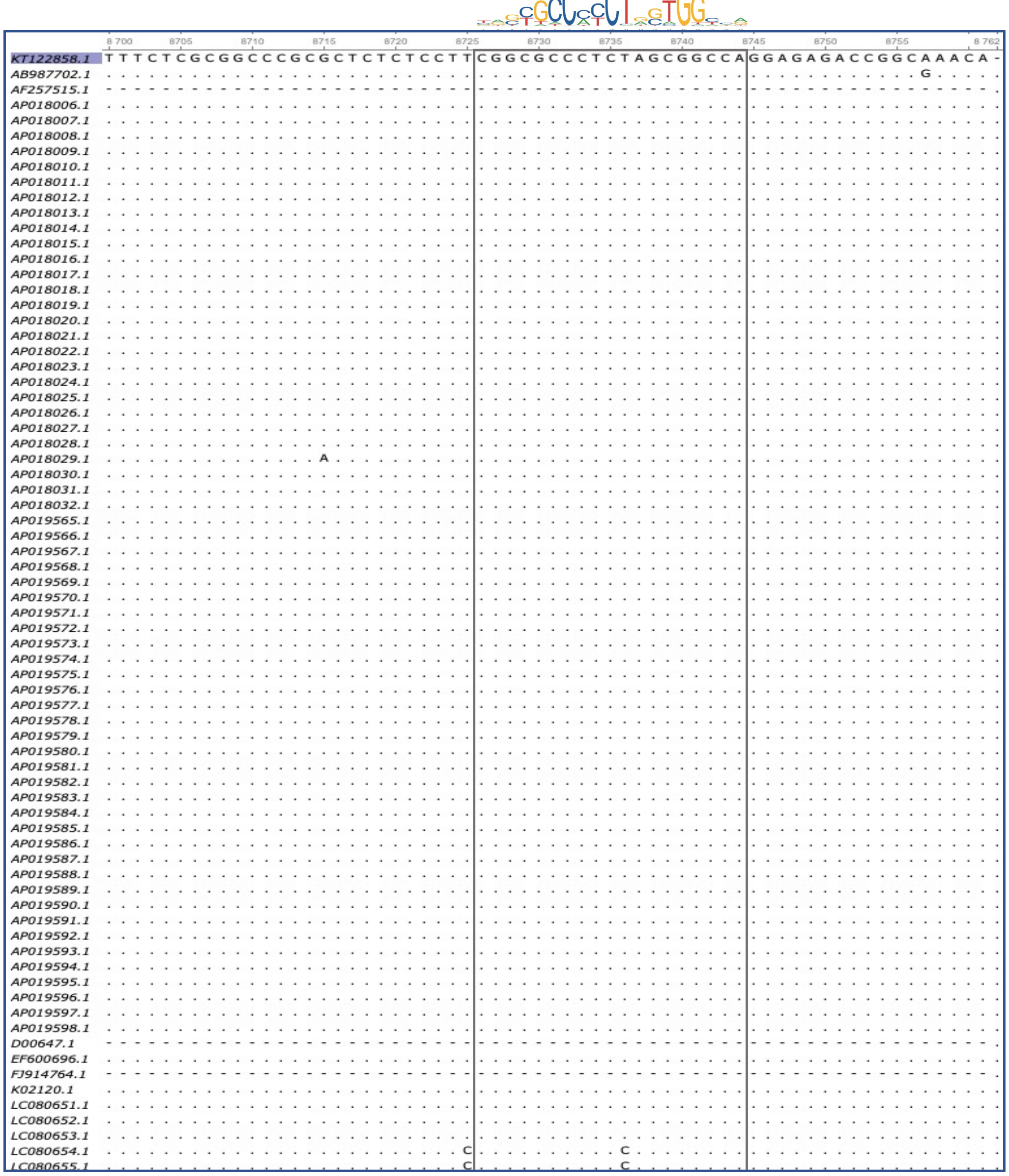

**Fig. S1 (continued).**

**3’ LTR**

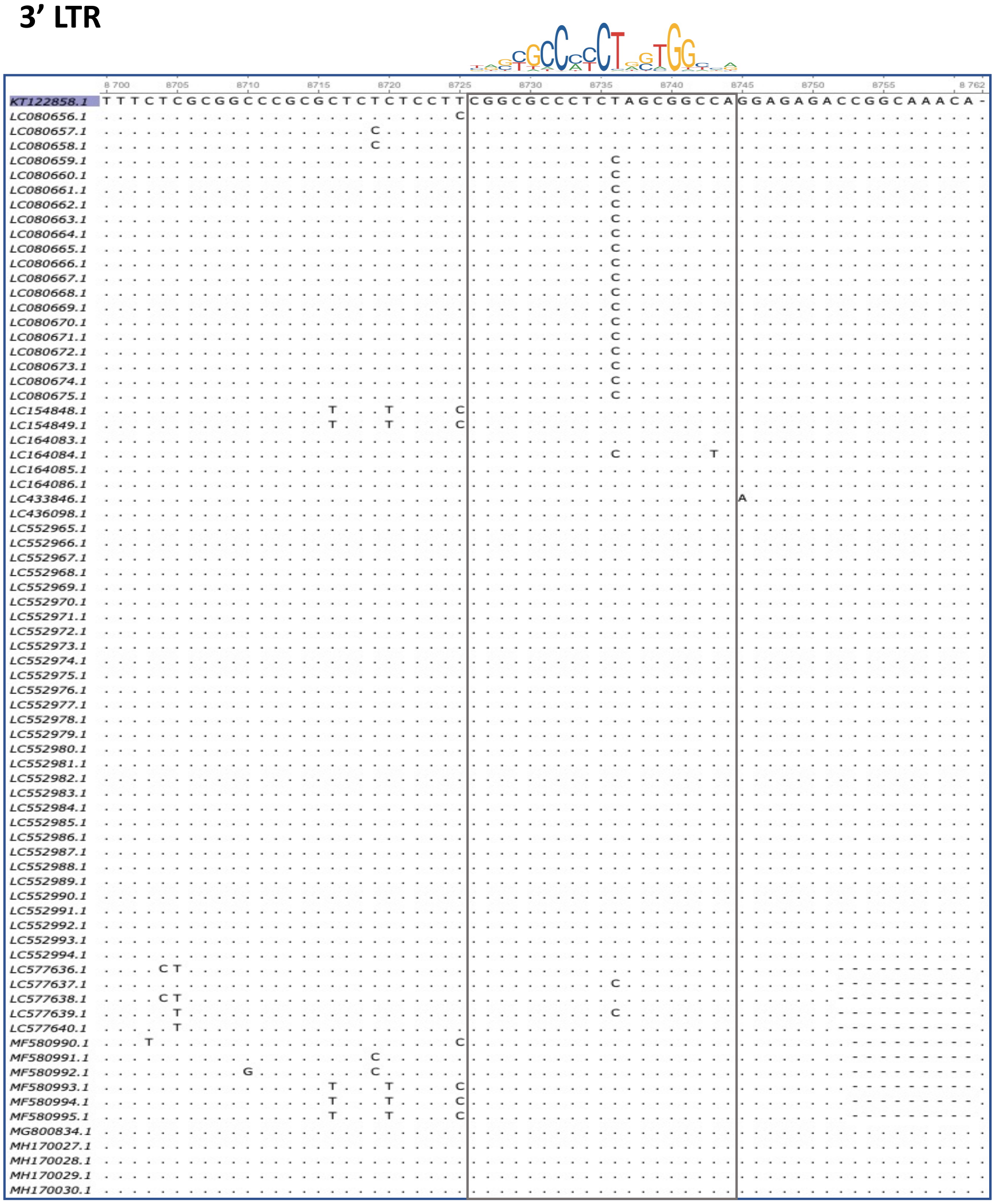

**Fig. S1 (continued).**

| Accession | PMID | Host |
|---|---|---|
| KT122858.1 | 27141823 | Ovis Aries |
| AB987702.1 | 28330779 | Bos Taurus |
| AF257515.1 | 11080485 | Bos Taurus |
| AP018006.1 | 29913249 | Bos Taurus |
| AP018007.1 | 29913249 | Bos Taurus |
| AP018008.1 | 29913249 | Bos Taurus |
| AP018009.1 | 29913249 | Bos Taurus |
| AP018010.1 | 29913249 | Bos Taurus |
| AP018011.1 | 29913249 | Bos Taurus |
| AP018012.1 | 29913249 | Bos Taurus |
| AP018013.1 | 29913249 | Bos Taurus |
| AP018014.1 | 29913249 | Bos Taurus |
| AP018015.1 | 29913249 | Bos Taurus |
| AP018016.1 | 29913249 | Bos Taurus |
| AP018017.1 | 29913249 | Bos Taurus |
| AP018018.1 | 29913249 | Bos Taurus |
| AP018019.1 | 29913249 | Bos Taurus |
| AP018020.1 | 29913249 | Bos Taurus |
| AP018021.1 | 29913249 | Bos Taurus |
| AP018022.1 | 29913249 | Bos Taurus |
| AP018023.1 | 29913249 | Bos Taurus |
| AP018024.1 | 29913249 | Bos Taurus |
| AP018025.1 | 29913249 | Bos Taurus |
| AP018026.1 | 29913249 | Bos Taurus |
| AP018027.1 | 29913249 | Bos Taurus |
| AP018028.1 | 29913249 | Bos Taurus |
| AP018029.1 | 29913249 | Bos Taurus |
| AP018030.1 | 29913249 | Bos Taurus |
| AP018031.1 | 29913249 | Bos Taurus |
| AP018032.1 | 29913249 | Bos Taurus |
| AP019565.1 | 29913249 | Bos Taurus |
| AP019566.1 | 29913249 | Bos Taurus |
| AP019567.1 | 29913249 | Bos Taurus |
| AP019568.1 | 29913249 | Bos Taurus |
| AP019569.1 | 29913249 | Bos Taurus |
| AP019570.1 | 29913249 | Bos Taurus |
| AP019571.1 | 29913249 | Bos Taurus |
| AP019572.1 | 29913249 | Bos Taurus |
| AP019573.1 | 29913249 | Bos Taurus |
| AP019574.1 | 29913249 | Bos Taurus |
| AP019575.1 | 29913249 | Bos Taurus |
| AP019576.1 | 29913249 | Bos Taurus |
| AP019577.1 | 29913249 | Bos Taurus |
| AP019578.1 | 29913249 | Bos Taurus |
| AP019579.1 | 29913249 | Bos Taurus |
| AP019580.1 | 29913249 | Bos Taurus |
| AP019581.1 | 29913249 | Bos Taurus |
| AP019582.1 | 29913249 | Bos Taurus |
| AP019583.1 | 29913249 | Bos Taurus |
| AP019584.1 | 29913249 | Bos Taurus |
| AP019585.1 | 29913249 | Bos Taurus |
| AP019586.1 | 29913249 | Bos Taurus |
| AP019587.1 | 29913249 | Bos Taurus |
| AP019588.1 | 29913249 | Bos Taurus |
| AP019589.1 | 29913249 | Bos Taurus |
| AP019590.1 | 29913249 | Bos Taurus |
| AP019591.1 | 29913249 | Bos Taurus |
| AP019592.1 | 29913249 | Bos Taurus |
| AP019593.1 | 29913249 | Bos Taurus |
| AP019594.1 | 29913249 | Bos Taurus |
| AP019595.1 | 29913249 | Bos Taurus |
| AP019596.1 | 29913249 | Bos Taurus |
| AP019597.1 | 29913249 | Bos Taurus |
| AP019598.1 | 29913249 | Bos Taurus |
| D00647.1 | 2167927 | Bos Taurus |
| EF600696.1 | 8230433 | Ovis Aries |
| FJ914764.1 | 19650931 | Bos Taurus |
| K02120.1 | 2983308 | Bos Taurus |
| LC080651.1 | 26754835 | Bos Taurus |
| LC080652.1 | 26754835 | Bos Taurus |
| LC080653.1 | 26754835 | Bos Taurus |
| LC080654.1 | 26754835 | Bos Taurus |
| LC080655.1 | 26754835 | Bos Taurus |
| KT122858.1 | 27141823 | Ovis Aries |

| Accession | PMID | Host |
|---|---|---|
| LC080656.1 | 26754835 | Bos Taurus |
| LC080657.1 | 26754835 | Bos Taurus |
| LC080658.1 | 26754835 | Bos Taurus |
| LC080659.1 | 26754835 | Bos Taurus |
| LC080660.1 | 26754835 | Bos Taurus |
| LC080661.1 | 26754835 | Bos Taurus |
| LC080662.1 | 26754835 | Bos Taurus |
| LC080663.1 | 26754835 | Bos Taurus |
| LC080664.1 | 26754835 | Bos Taurus |
| LC080665.1 | 26754835 | Bos Taurus |
| LC080666.1 | 26754835 | Bos Taurus |
| LC080667.1 | 26754835 | Bos Taurus |
| LC080668.1 | 26754835 | Bos Taurus |
| LC080669.1 | 26754835 | Bos Taurus |
| LC080670.1 | 26754835 | Bos Taurus |
| LC080671.1 | 26754835 | Bos Taurus |
| LC080672.1 | 26754835 | Bos Taurus |
| LC080673.1 | 26754835 | Bos Taurus |
| LC080674.1 | 26754835 | Bos Taurus |
| LC080675.1 | 26754835 | Bos Taurus |
| LC154848.1 | 27771791 | Bos Taurus |
| LC154849.1 | 27771791 | Bos Taurus |
| LC164083.1 | 27534623 | Bos Taurus |
| LC164084.1 | 27534623 | Bos Taurus |
| LC164085.1 | 27534623 | Bos Taurus |
| LC164086.1 | 27534623 | Bos Taurus |
| LC433846.1 | 32122602 | Bos Taurus |
| LC436098.1 | 32122602 | Bos Taurus |
| LC552965.1 | 33486630 | Bos Taurus |
| LC552966.1 | 33486630 | Bos Taurus |
| LC552967.1 | 33486630 | Bos Taurus |
| LC552968.1 | 33486630 | Bos Taurus |
| LC552969.1 | 33486630 | Bos Taurus |
| LC552970.1 | 33486630 | Bos Taurus |
| LC552971.1 | 33486630 | Bos Taurus |
| LC552972.1 | 33486630 | Bos Taurus |
| LC552973.1 | 33486630 | Bos Taurus |
| LC552974.1 | 33486630 | Bos Taurus |
| LC552975.1 | 33486630 | Bos Taurus |
| LC552976.1 | 33486630 | Bos Taurus |
| LC552977.1 | 33486630 | Bos Taurus |
| LC552978.1 | 33486630 | Bos Taurus |
| LC552979.1 | 33486630 | Bos Taurus |
| LC552980.1 | 33486630 | Bos Taurus |
| LC552981.1 | 33486630 | Bos Taurus |
| LC552982.1 | 33486630 | Bos Taurus |
| LC552983.1 | 33486630 | Bos Taurus |
| LC552984.1 | 33486630 | Bos Taurus |
| LC552985.1 | 33486630 | Bos Taurus |
| LC552986.1 | 33486630 | Bos Taurus |
| LC552987.1 | 33486630 | Bos Taurus |
| LC552988.1 | 33486630 | Bos Taurus |
| LC552989.1 | 33486630 | Bos Taurus |
| LC552990.1 | 33486630 | Bos Taurus |
| LC552991.1 | 33486630 | Bos Taurus |
| LC552992.1 | 33486630 | Bos Taurus |
| LC552993.1 | 33486630 | Bos Taurus |
| LC552994.1 | 33486630 | Bos Taurus |
| LC577636.1 | 33633166 | Bos Taurus |
| LC577637.1 | 33633166 | Bos Taurus |
| LC577638.1 | 33633166 | Bos Taurus |
| LC577639.1 | 33633166 | Bos Taurus |
| LC577640.1 | 33633166 | Bos Taurus |
| MF580990.1 | 29224130 | Bos grunniens |
| MF580991.1 | 29224130 | Bos grunniens |
| MF580992.1 | 29224130 | Bos grunniens |
| MF580993.1 | 29224130 | Bos grunniens |
| MF580994.1 | 29224130 | Bos grunniens |
| MF580995.1 | 29224130 | Bos grunniens |
| MG800834.1 | 31142319 | Bos Taurus |
| MH170027.1 | 30039314 | Bos Taurus |
| MH170028.1 | 30039314 | Bos Taurus |
| MH170029.1 | 30039314 | Bos Taurus |
| MH170030.1 | 30039314 | Bos Taurus |

**Table S1: List of sequences used in Fig. S1.**

| | | |
|---|---|---|
| **Cloning** | | |
| Mutagenesis-LTRmCTCF-fw | CV4203 | CTCCTTCGGCGCCCTTAAGATATCAGGAGAG |
| Mutagenesis-LTRmCTCF-rv | CV4204 | CTCTCCTGATATCTTAAGGGCGCCGAAGGAG |
| Lenti-LTR-BLV-AgeI-fw | CV4387 | ATATTCTAGAGCTCGAGATCGGGTGTATG |
| Lenti-LTR-BLV-XbaI-rv | CV4388 | ATATACCGGTGCCAAGCTTACTTAGATCG |
| **ChIP-qPCR** | | |
| Lenti-LTR-fw | CV3081 | GCTCTCTCCTTCGGCGCCCT |
| Lenti-LTR-rv | CV4425 | GCGACCGGTGCCAAGCTTAC |
| Positive controle CTCF (MDM2)-fw | CV4427 | TGTATGAACGCATACCTGCCC |
| Positive controle CTCF (MDM2)-rv | CV4428 | CATCATGCCATCTAGCGGTCT |
| Negative contrôle CTCF (GAPDH TATA)-fw | CV2004 | GCCCCCGGTTTCTATAAATTG |
| Negative contrôle CTCF (GAPDH TATA)-rv | CV2005 | AGAAGATGCGGCTGACTGTC |
| Rasa3 E1-fw | CV3168 | GACCCAGCATGGCGGTGGAG |
| Rasa3 E1-rv | CV3169 | CCGCCTGCTCCTGCGAACTC |
| Rasa3 I1-fw | CV3170 | CATACCTCCCTGGCCCCGCT |
| Rasa3 I1-rv | CV3171 | CAGAAGCACCCGCGCTCCAA |
| 5’ junction (L267)-fw | CV3174 | CCGGCAGCTTCTGACCGCAG |
| 5' junction (L267)-rv | CV3175 | CGCCTAGGCCGGCATGATCT |
| 5' junction (YR2)-fw | CV3617 | TTCTCTAATTCTCCACTTTCCCAA |
| 5' junction (YR2)-rv | CV3618 | CCTAGGCCGGCATGATCTTT |
| 5' junction (M2241)-fw | CV3661 | TCCACCCATAGTTCATCAGGT |
| 5' junction (M2241)-rv | CV3662 | CTAGGCCGGCATGATCTTTC |
| LTR5’-fw | CV3123 | AAGGGCGTCTGGCTTGCACC |
| LTR5’-rv | CV3124 | AATCCCGGACGAGCCCCCAA |
| gag-fw | CV3028 | AGCCCAACGCCGGGGATCTT |
| gag-rv | CV3029 | CGGGGCCTTGGACGATGGTG |
| pro-fw | CV3030 | TTCCTCTGGCTCGCAGCCGT |
| pro-rv | CV3031 | TTCAGCCCCGGTGTCCACGA |
| pol-fw | CV3032 | TTCCTGCGGCCCTTTGCCTC |
| pol-rv | CV3033 | AGCCCGCCAAGAGACCTGCT |
| Tax/Rex E1-fw | CV3034 | TGGCTAGGACCACTCCCGGC |
| Tax/Rex E1-rv | CV3035 | CGGTTGTGGGCGTCTTCGGG |
| env-fw | CV3038 | CCAGAACCGACGGGGGCTTG |
| env-rv | CV3039 | GCTGGAGATCACCGAGGCGG |
| miRNAs-fw | CV3127 | ACGCCCTGTTGCACACCCTT |
| miRNAs-rv | CV3128 | CTCAGAACCCGGGGCCTTGC |
| Tax/Rex E2-fw | CV3044 | CTTGTGGACCCCCTCCGGCT |
| Tax/Rex E2-rv | CV3045 | AGGGCTCGCCTAGGGGTAGAA |
| LTR3’ (L267 and YR2)-fw | CV3046 | TGGTTGCTAGCGGGAAACTAAGACT |
| LTR3’ (L267, YR2 and M2241)-rv | CV3047 | CTGGTTTACGGGGCGGTGGC |
| LTR3' (M2241)-fw | CV4448 | AGAAAATGAATGGCTCTCCCGCCT |
| 3’ junction (L267)-fw | CV3176 | AAGGGCGTCTGGCTTGCACC |
| 3’ junction (L267)-rv | CV3177 | GGCCGTCGAGTTCCGACCTG |
| 3’ junction (YR2)-fw | CV3130 | ACTTTCTGTTTCTCGCGGCCC |
| 3’ junction (YR2)-rv | CV3614 | GAGTTTAAAATATTCTCCCTTATCATGTAC |
| 3’ junction (M2241)-fw | CV3664 | TCTAGCGGCCAGGAGAGA |
| 3’ junction (M2241)-rv | CV3679 | TGATCTGCCCAAATTGCCAA |
| Rasa3 I2-fw | CV3179 | TCCCGAGCCCTAGGTGTGCC |
| Rasa3 I2-rv | CV3180 | CTCCTCTCCCGCAGGCCAGT |
| Rasa3 E2-fw | CV3181 | TCCCACATACCCGGGGCCAA |
| Rasa3 E2-rv | CV3182 | GGTCCTGAAAACCTCCTCCTGGT |
| **4C-seq** | | |
| **Divergent primers** | | |
| VP1 (L267 and YR2)-fw | CV4295 | TACACGACGCTCTTCCGATCTAAACAATTGGGGGCTCGT |
| VP1 (L267 and YR2)-rv | CV4296 | ACTGGAGTTCAGACGTGTGCTCTTCCGATCTTTTTATCAGCAGGTGAGGTC |
| VP2 L267-fw | CV4301 | TACACGACGCTCTTCCGATCTCCTTCGAGCTCTTCGGGATC |
| VP2 L267-rv | CV4302 | ACTGGAGTTCAGACGTGTGCTCTTCCGATCTTCAGTCATGCACTCAGAGAG |
| VP2 YR2-fw | CV4305 | TACACGACGCTCTTCCGATCTCCTTCAAGCTCTTCGGGATC |
| VP2 YR2-rv | CV4302 | ACTGGAGTTCAGACGTGTGCTCTTCCGATCTTCAGTCATGCACTCAGAGAG |
| VP3 L267-fw | CV4304 | TACACGACGCTCTTCCGATCTGGAACTCGACGGCCAAGATC |
| VP3 L267-rv | CV4296 | ACTGGAGTTCAGACGTGTGCTCTTCCGATCTTTTTATCAGCAGGTGAGGTC |
| VP3 YR2-fw | CV4310 | TACACGACGCTCTTCCGATCTGCTGCCTATAGTTTTAGAGCTT |
| VP3 YR2-rv | CV4311 | ACTGGAGTTCAGACGTGTGCTCTTCCGATCTCCCATTTTTAGTCTGTAAGGA |
| **Sequencing library preparation** | | |
| L267 rep 1#-fw | CV4328 | AATGATACGGCGACCACCGAGATCTACACTCTTTCCCTACACGACGCTCTTCCGATCT |
| L267 rep 1#-rv | CV4331 | CAAGCAGAAGACGGCATACGAGATCACTGTGTGACTGGAGTTCAGACGTGTGCT |
| L267 rep 2#-fw | CV4328 | AATGATACGGCGACCACCGAGATCTACACTCTTTCCCTACACGACGCTCTTCCGATCT |
| L267 rep 2#-rv | CV4334 | CAAGCAGAAGACGGCATACGAGATTACAAGGTGACTGGAGTTCAGACGTGTGCT |
| YR2 rep 1#-fw | CV4328 | AATGATACGGCGACCACCGAGATCTACACTCTTTCCCTACACGACGCTCTTCCGATCT |
| YR2 rep 1#-rv | CV4332 | CAAGCAGAAGACGGCATACGAGATATTGGCGTGACTGGAGTTCAGACGTGTGCT |
| YR2 rep 2#-fw | CV4328 | AATGATACGGCGACCACCGAGATCTACACTCTTTCCCTACACGACGCTCTTCCGATCT |
| YR2 rep 2#-rv | CV4336 | CAAGCAGAAGACGGCATACGAGATCGTTTCACGTGACTGGAGTTCAGACGTGTGCT |

**Table S2: List of oligonucleotides used in this study.**

| | | |
|---|---|---|
| IgG | C15410206 | Diagenode |
| PanH3 | 07-690 | Millipore |
| AcH3 | Ab47915 | Abcam |
| H3K4me2 | 07-030 | Millipore |
| H3K4me3 | 04-745 | Millipore |
| H3K9Ac | C15410004 | Diagenode |
| H3K27Ac | Ab4729 | Abcam |
| CTCF | 07-729 | Millipore |
| Rad21 | Ab992 | Abcam |

**Table S3: List of antibodies used in this study.**

| Sample Name | Integration Site | Impacted host gene | Provirus orientation relative to reference genome |
|---|---|---|---|
| L267 | chr10:86299813 | RASA3 | - |
| YR2 | chr17:18211332 | ELF2 | - |
| M2241 | chr5:20532551 | RAPGEF6 | + |

Reference BLV genome: KT122858.1 (Genbank)
Reference ovine genome: Oar_v3.1 (UCSC)

**Table S4: List of BLV integration sites.**

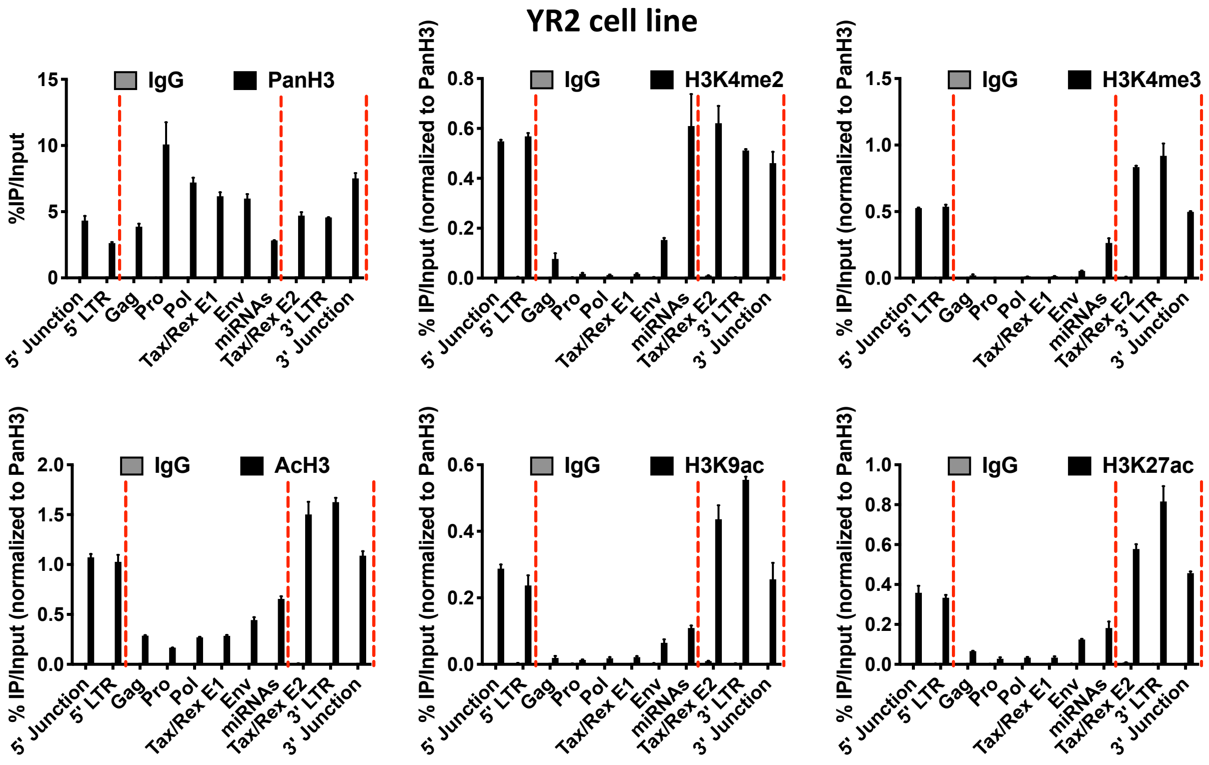


**Fig. S2**: **CTCF defines a specific epigenetic profile along the BLV proviral genome.** Chromatin prepared from YR2 cells was immunoprecipitated with specific antibodies directed against histone H3 (PanH3), different histone post-translational modifications (H3Kme2, H3Kme3, AcH3, H3K9ac, H3K27ac, H3k27me3, H3K36me3) or with an IgG as background measurement. Purified DNA was then amplified with oligonucleotide primers hybridizing to the BLV proviral genome. Results are presented as histograms indicating percentages of immunoprecipitated DNA compared to the input DNA (% IP/Input) normalized to PanH3. Data are the means ± SD from one representative of at least three independent experiments.

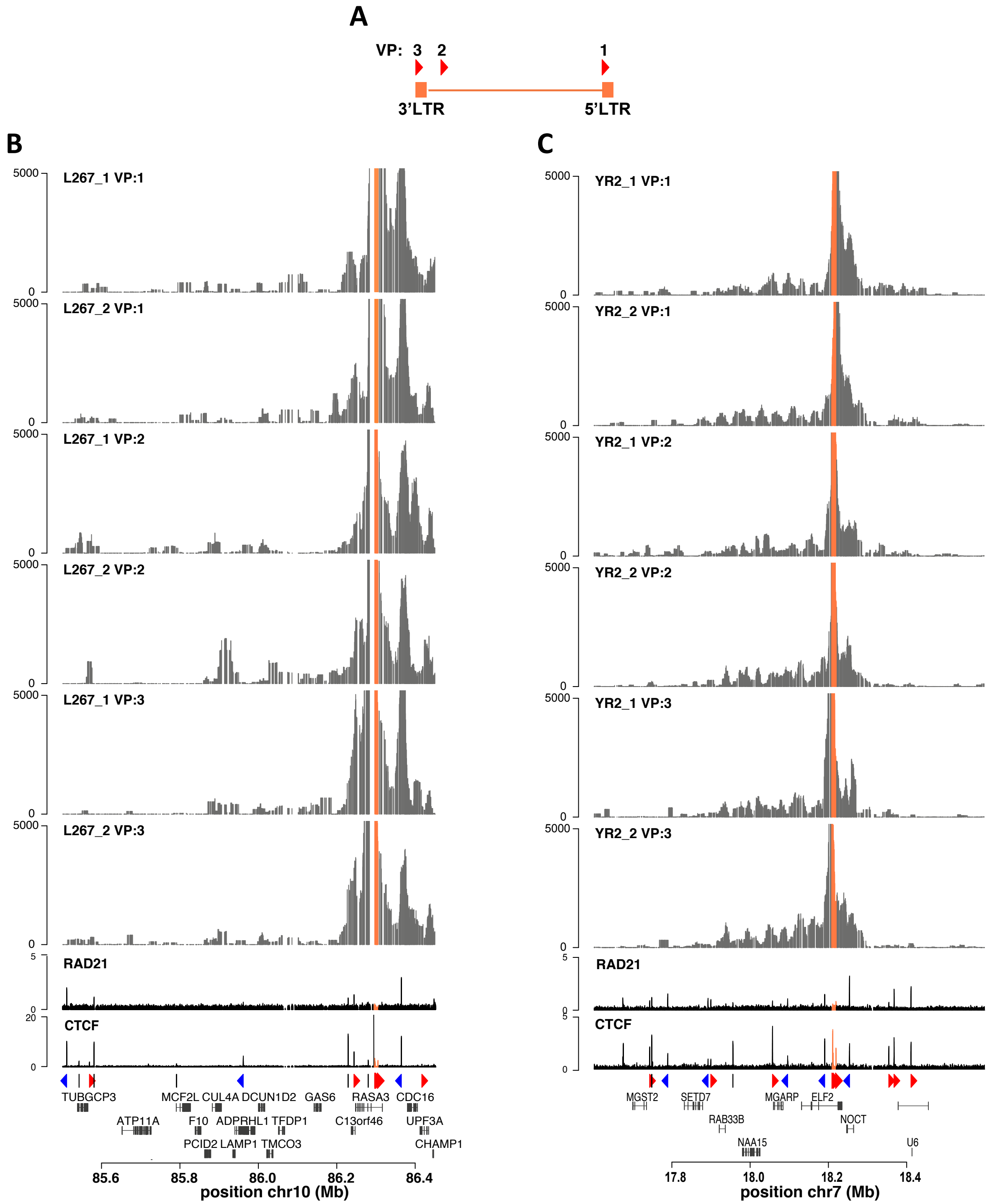


**Fig. S3: Biological replicates of 4C-seq profiles using 3 different viewpoints. (A)** Localization of the 3 viewpoints (VP) in the BLV provirus. **(B,C)** Replicates of 4C-seq contact profiles in the context of L267 cells or YR2 cells using each viral CTCF binding sites as viewpoint. Below the 4C plots are shown the Rad21 and CTCF ChIP-seq profiles. The CTCF binding motifs orientation is indicated by red (forward) or blue (reverse) arrows or by gray bars (undetermined) as well as the position of host cellular genes, with respect to the previously identified BLV integration site.
